# Thrombolysis exacerbates cerebrovascular injury after ischemic stroke via a VEGF-B dependent effect on adipose lipolysis

**DOI:** 10.1101/2024.10.11.617532

**Authors:** Ingrid Nilsson, Enming J. Su, Linda Fredriksson, Benjamin Heller Sahlgren, Zsuzsa Bagoly, Christine Moessinger, Christina Stefanitsch, Frank Chenfei Ning, Manuel Zeitelhofer, Lars Muhl, Anna-Lisa E. Lawrence, Pierre D. Scotney, Li Lu, Erik Samén, Heidi Ho, Richard F. Keep, Robert L. Medcalf, Daniel A. Lawrence, Ulf Eriksson

## Abstract

Cerebrovascular injuries leading to edema and hemorrhage after ischemic stroke are common. The mechanisms underlying these events and how they are connected to known risk factors for poor outcome, like obesity and diabetes, is relatively unknown. Herein we demonstrate that increased adipose tissue lipolysis is a dominating risk factor for the development of a compromised cerebrovasculature in ischemic stroke. Reducing adipose lipolysis by VEGF-B antagonism improved vascular integrity by reducing ectopic cerebrovascular lipid deposition. Thrombolytic therapy in ischemic stroke using tissue plasminogen activator (tPA) leads to increased risk of hemorrhagic complications, substantially limiting the use of thrombolytic therapy. We provide evidence that thrombolysis with tPA promotes adipose tissue lipolysis, leading to a rise in plasma fatty acids and lipid accumulation in the ischemic cerebrovasculature after stroke. VEGF-B blockade improved the efficacy and safety of thrombolysis suggesting the potential use of anti-VEGF-B therapy to extend the therapeutic window for stroke management.

## Introduction

Obesity, type-2-diabetes and other inter-related metabolic risk factors for cardiovascular diseases, including acute ischemic stroke, comprise the metabolic syndrome.^1^ Overt stroke phenotypes associated with the metabolic syndrome include augmented thrombosis, edema, intracerebral hemorrhage (ICH) and pro-inflammatory responses, correlating with poor functional outcome.^2–5^ Admission hyperglycemia has furthermore been identified as an independent risk factor for poor outcomes after acute ischemic stroke, including increased mortality, larger infarct size, and increased risk of ICH.^2,6–10^ However, intensive blood glucose management after acute ischemic stroke does not improve outcome or reduce the incidence of further strokes.^11–13^ The pathophysiological mechanisms underlying the connection between the metabolic syndrome and poor outcome after acute ischemic stroke thus remain poorly understood.^14^

Insulin resistance and excessive fatty acid (FA) flux are hallmarks of the metabolic syndrome.^1^ Vascular endothelial growth factor-B (VEGF-B) signaling has been shown to promote FA flux and is highly expressed in metabolically active tissues such as heart and skeletal muscle.^15–18^ VEGF-B downstream signaling requires engagement with both VEGF receptor-1 (VEGFR1) and neuropilin-1^19,20^, co-expressed on vascular endothelial cells (ECs), resulting in increased trans-endothelial long-chain FA transport.^16^ Inhibiting overall VEGF-B signaling consequently leads to decreased FA uptake and lipid droplet (LD) accumulation in heart and skeletal muscle, but also decreased circulating levels of non-esterified FAs (NEFAs) and triglycerides (TGs), correlating with improved insulin sensitivity in experimental models of obesity and type-2-diabetes.^16,21,22^ More recently, we demonstrated that limiting VEGF-B signaling specifically in white adipose tissue (WAT) is sufficient to restore insulin sensitivity by inhibiting FA flux from visceral adipose tissue to blood, a process termed lipolysis.^23^

Intravenous thrombolysis with recombinant tissue plasminogen activator (rtPA): alteplase, or more recently, tenecteplase, is the only approved pharmacological treatment for acute ischemic stroke. The efficacy of thrombolysis is demonstrated up to 4.5 h after stroke symptom onset^24,25^, however, the benefit is limited by an increased risk of edema and ICH.^26,27^ Furthermore, incomplete recanalization and increased bleeding risk is observed after thrombolysis in people exhibiting features of the metabolic syndrome.^2,5,8,10,27^ Edema and ICH after ischemic stroke result from compromised blood-brain barrier (BBB) integrity.^27–30^ The integrity of the BBB is maintained by both EC tight junctions, limiting paracellular permeability, and restrained vesicular trafficking, limiting transcellular permeability^31^. Breach of the BBB and the underlying basal lamina can result in ICH after reperfusion.^27,31,32^ Thus, preserving BBB integrity during thrombolysis can reduce the risk of symptomatic ICH (sICH), a potentially fatal adverse event after ischemic stroke.

Herein, we present evidence of a previously unrecognized acute effect of thrombolysis, involving tPA-mediated activation of WAT lipolysis and subsequent release of NEFAs into the blood circulation. Concomitant buildup of LDs in the ischemic cerebrovasculature correlated with BBB disruption and ICH incidence. Limiting WAT lipolysis through pharmacological inhibition of VEGF-B in conjunction with thrombolysis resulted in attenuated ICH and improved outcome after ischemia, signifying a new strategy to expand the therapeutic window for thrombolysis with rtPA.

## Results

### A fatty acid-rich environment promotes fatty acid uptake and impairs glucose uptake and integrity in endothelial cells

To model effects of obesity and the metabolic syndrome on ECs, we exposed primary human umbilical vein ECs (HUVECs) to high long-chain FA (^hi^FA) and/or glucose (^hi^Glc) levels for 2 h, 24 h and five days (**Figure 1A**). Co-exposure decreased glucose uptake at all time-points (**Figure 1B**). ^hi^Glc alone, mimicking hyperglycemia, did not influence glucose uptake, whereas FA exposure, also in moderate concentrations, decreased glucose uptake capacity (**Figures 1B and 1C**), indicating that a FA-rich environment impairs EC glucose uptake *in vitro*. ^hi^FA exposure down-regulated glucose transporter 1 (GLUT1) distribution (**Figure 1D and 1E**), suggesting a shift in nutrient uptake specificity. Indeed, HUVECs incubated in FA-rich media conversely increased long-chain FA uptake (**Figure 1F**), correlating with augmented mitochondria activity (**Figure 1G**) and induction of neutral lipid synthesis and storage in adipophilin/perilipin-2 (PLIN2+) intracellular LDs (**Figures 1H, 1I, 1J, S1A** and **S1B**). Increased long-chain FA uptake correlated with weakened paracellular integrity and increased macro-pinocytosis (**Figures 1K, 1L, 1M and 1N**), implying that FA uptake compromises barrier integrity. Taken together, our data demonstrates that long-chain FA exposure will switch ECs from glucose to FA uptake, coinciding with declined EC barrier properties.

**Figure 1.**
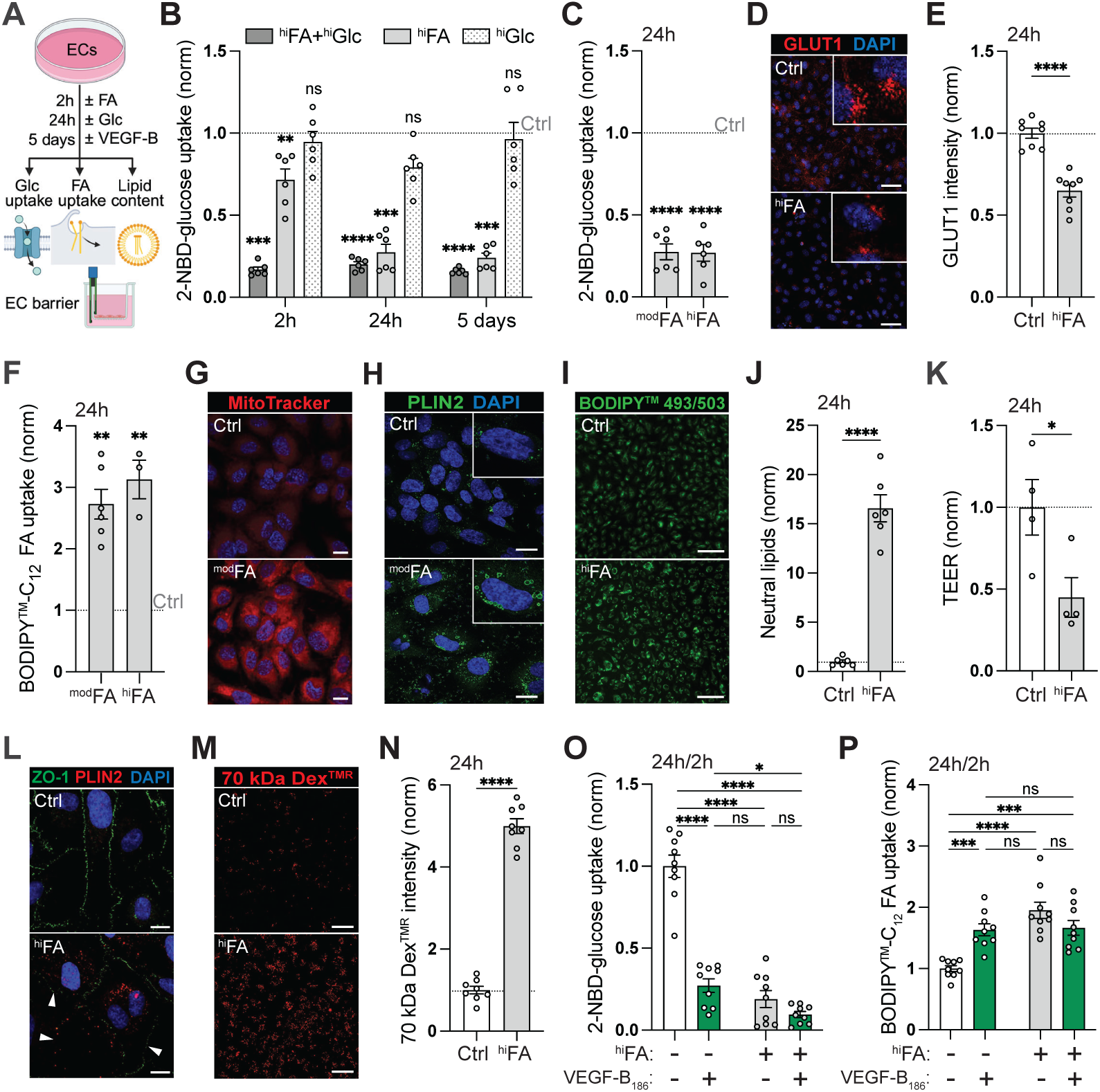
A fatty acid-rich environment promotes fatty acid uptake and impairs glucose uptake and integrity in endothelial cells. **(A)** Schematic representation of the *in vitro* experimental outline in HUVECs. **(B)** Glucose uptake in response to high long-chain FA (^hi^FA; 0.1 mM, oleic/palmitic acid mix) and/or glucose (^hi^Glc, 20 mM D-Glucose) levels, compared to cells cultured in control media (Ctrl) for the indicated timepoints. **(C)** Glucose uptake after 0.05 mM (^mod^FA) or 0.5 mM (^hi^FA) oleic acid. **(D-E)** GLUT1 intensity after ^hi^FA exposure. Insets show higher magnification. **(F)** FA uptake after ^mod^FA or ^hi^FA exposure. **(G-H)** Mitochondria (G) and PLIN2+ LD (H) visualization after ^mod^FA exposure (0.05 mM; oleic/palmitic acid mix). Insets show higher magnification. **(I-J)** Neutral lipid content after ^hi^FA exposure (0.5 mM oleic acid). **(K)** Trans-endothelial electrical resistance after ^hi^FA exposure. **(L)** ZO-1 intensity (arrowheads) after ^hi^FA exposure. LDs stained with PLIN2. **(M-N)** Dex^TMR^ uptake after ^hi^FA exposure. **(O-P)** Glucose (O) and FA (P) uptake after VEGF-B stimulation in the absence or presence of ^hi^FAs exposure. Data presented as mean ± s.e.m of 3-9 biological replicates per condition, normalized to control treated cells. Scale bars, 10 μm (D, G, H, L); 50 μm (I, M). Statistical evaluation using unpaired *t*-test with Welch’s correction (B, F, J), unpaired *t*-test (C, E, K, N), or two-way ANOVA followed by Tukeýs multiple comparisons test (O-P). \**P*<0.05, \*\**P*<0.01, \*\*\**P*<0.001, \*\*\*\**P*<0.0001. See also Figure S1.

Analogous to excess FA supplementation, VEGF-B stimulation induces a shift from glucose to FA uptake in various types of ECs.^16,33^ No additive or synergistic effect was observed when stimulating HUVECs with VEGF-B on top of ^hi^FA treatment (**Figures 1O and 1P**), suggesting a shared mechanism underlying increased FA uptake through VEGF-B signaling and excess FA exposure, and the impairment of glucose uptake. To examine the generalizability of these findings to brain endothelium, select experiments were repeated in primary brain microvascular ECs (HBMECs). HBMECs phenocopied the shift from glucose to FA uptake in response to VEGF-B, whereas a neutralizing anti-VEGF-B (αVEGF-B) monoclonal antibody had no effect, supporting a paracrine and not autocrine source of VEGF-B (**Figures S1C, S1D, S1E, S1F, S1G and S1H**). *Vegfb* is widely expressed in brain, with the most distinct expression in neurons and ependymal cells. *Vegfb* expressing cells were also found around arteriolar-sized blood vessels (**Figure S2A**). VEGFR1 and neuropilin-1 were co-expressed specifically in ECs, in agreement with a paracrine signaling modality where VEGF-B is secreted from neighboring parenchymal cells or circulating in the blood **(Figures S2B and** S2C).^16,17,23^

### Short-term inhibition of VEGF-B signaling in high-fat diet mice lowers circulating fatty acid and triglyceride levels, restoring brain glucose uptake capacity

High-fat diet (HFD) mice develop obesity and characteristics of the metabolic syndrome, including overt weight gain and hyperglycemia (**Figures S3A, S3B and S3C**). Long-term treatment with αVEGF-B antibodies in HFD mice reduces plasma NEFA and TG levels by limiting FA transport from WAT to blood.^21,23^ Short-term treatment with αVEGF-B antibodies (one week) was sufficient to reproduce this effect in HFD mice (**Figures 2A, 2B and 2C**). In addition to lowering plasma NEFAs and TGs, αVEGF-B treatment decreased VLDL/LDL-cholesterol, whereas HDL-cholesterol and ketone bodies were unchanged (**Figures 2D, 2E and 2F**). Age-matched littermate mice maintained on grain-based chow diet did not exhibit significant changes after αVEGF-B treatment. Blood glucose and body weight remained unaffected (**Figures 2G and 2H**), suggesting that short-term αVEGF-B treatment has a selective effect on circulating lipids in cases of high-fat feeding or excess caloric intake, without impacting overall energy homeostasis.

**Figure 2.**
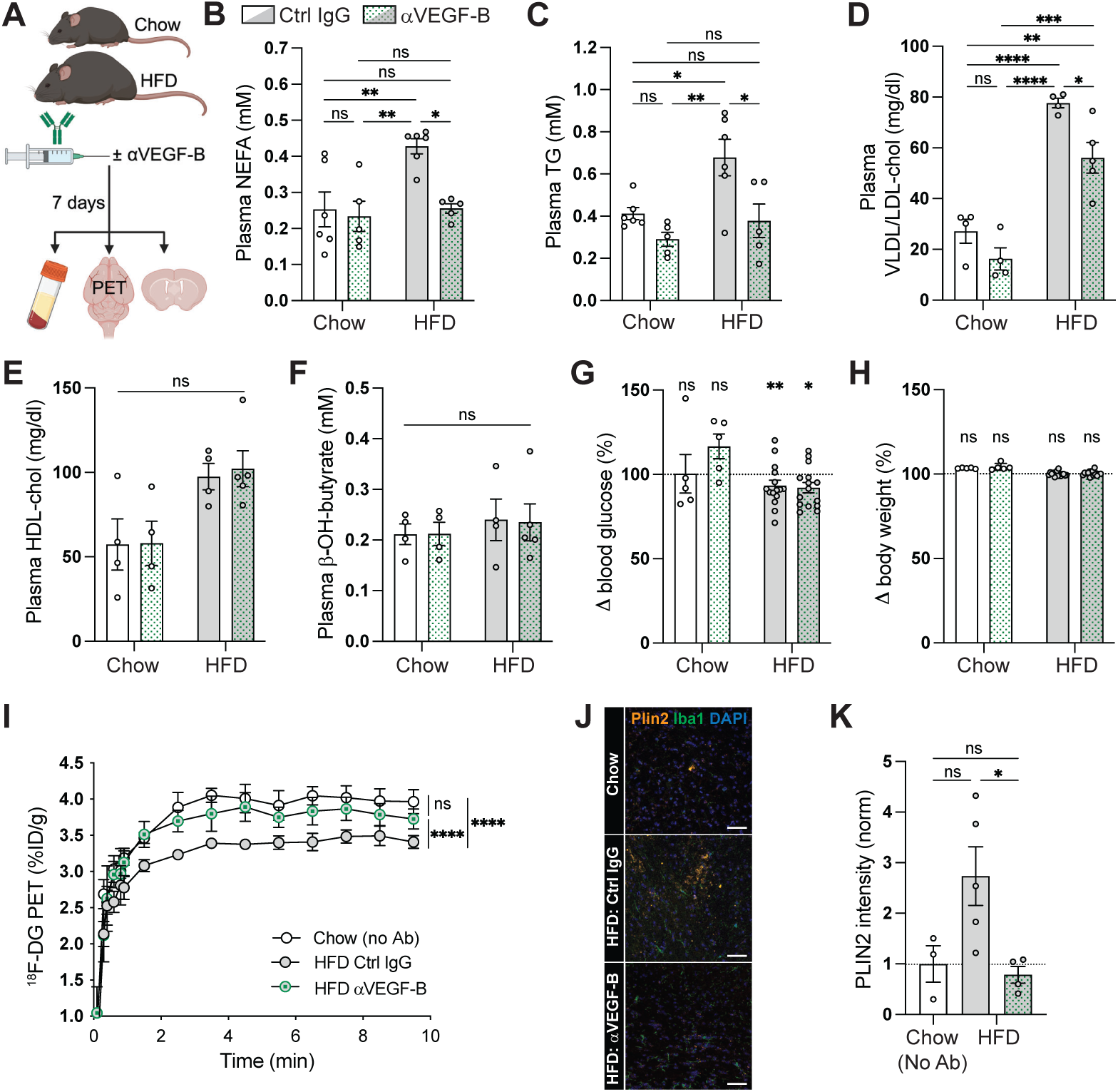
Short-term inhibition of VEGF-B signaling in high-fat diet mice lowers circulating fatty acid and triglyceride levels, restoring brain glucose uptake capacity. **(A)** Schematic representation of experimental outline. **(B-H)** Plasma NEFAs (B), plasma TGs (C), plasma VLDL/LDL cholesterol (D), plasma HDL cholesterol (E), blood ketones (F), blood glucose change (G) and body weight change (H) after short-term (one week) systemic αVEGF-B or isotype control IgG treatment of chow and HFD mice. **(I-K)** Dynamic brain glucose uptake (I) and LD accumulation (J-K) in treated HFD mice compared to non-treated age-matched chow mice. Scale bars, 50 μm. Data presented as mean ± s.e.m. based on n=5-6 mice/group (B-C), n=4-5 mice/group (D-F), n=5+15 (chow *vs.* HFD) mice/group (G-H), n=3-5 mice/group (I), and n=3-5 mice/group (K). Statistical evaluation using two-way ANOVA followed by Tukeýs multiple comparisons test (B-F, I), Wilcoxon matched-pairs signed rank test (pre *vs.* post, G-H) or one-way ANOVA followed by Tukeýs multiple comparisons test (K). \**P*<0.05, \*\**P*<0.01, \*\*\**P*<0.001, \*\*\*\**P*<0.0001. See also Figure S2 and S3.

HFD mice displayed significantly decreased brain glucose uptake capacity (**Figure 2I**). mRNA expression of *Slc2a1* (Glut1), the main and rate-limiting glucose transporter in brain, was however not significantly changed (**Figure S3D**), implying post-transcriptional regulation of brain glucose uptake in HFD mice. In addition, slight impairment of cerebrovascular integrity was observed (**Figure S3E**). Global *Vegfb* mRNA expression was however not changed (**Figure S3F**), arguing against local upregulation of brain VEGF-B signaling in HFD mice under physiological circumstances. Adipophilin/Plin2 is a marker for neutral lipid accumulation in metabolically active cells such as skeletal myocytes (**Figure S3G**) and diseases characterized with metabolic dysregulation.^34^ In healthy mouse brain, Plin2+ LDs were found primarily inside ventricular ependymal cells, and otherwise at low presence (**Figure S3H**), consistent with high metabolic activity in ependymal cells and low overall uptake and storage of neutral lipids in brain. Brain LD abundance was reduced, and glucose uptake capacity restored, after αVEGF-B treatment of HFD mice (**Figures 2I, 2J and 2K**), in agreement with a shift towards glucose utilization. These *in vivo* data thus confirm and extend the *in vitro* observations; suggesting that impairment of brain glucose uptake capacity in HFD mice is primarily regulated by increased circulating NEFAs and not by local *Vegfb* expression.

### High-fat diet feeding confers a pro-thrombotic and reperfusion-resistant phenotype after acute ischemic stroke, characterized by vascular hyperpermeability and ectopic endothelial lipid droplet accumulation

There are several determinants of outcome after acute ischemic stroke, including thrombogenicity, reperfusion efficiency, and BBB integrity linked to edema and ICH.^27,35^ The photo-thrombotic middle cerebral artery occlusion (MCAO) model^30^ was used to induce ischemic stroke in chow and HFD mice (**Figure 3A**). Infarct volume increased by 35% in HFD mice (**Figure 3B**), and an increased incidence and magnitude of ICH was observed (**Figure 3C**), consistent with previous reports.^36,37^ To determine if HFD mice were more prone to thrombosis, a separate cohort of mice were subjected to a milder photothrombotic insult. Measurement of cerebral blood flow (CBF) revealed a more pronounced initial CBF decline in HFD mice, as well as refractory spontaneous reperfusion, compared to chow-fed mice (**Figure 3D**). Furthermore, intravenous injection of a fibrinogen tracer right before MCAO induction revealed more pronounced vascular fibrin deposition (**Figures 3E, 3F and 3G**), collectively illustrating that HFD mice display a pro-thrombogenic and reperfusion-resistant stroke phenotype that presumably accounts for the increased infarct volume.

**Figure 3.**
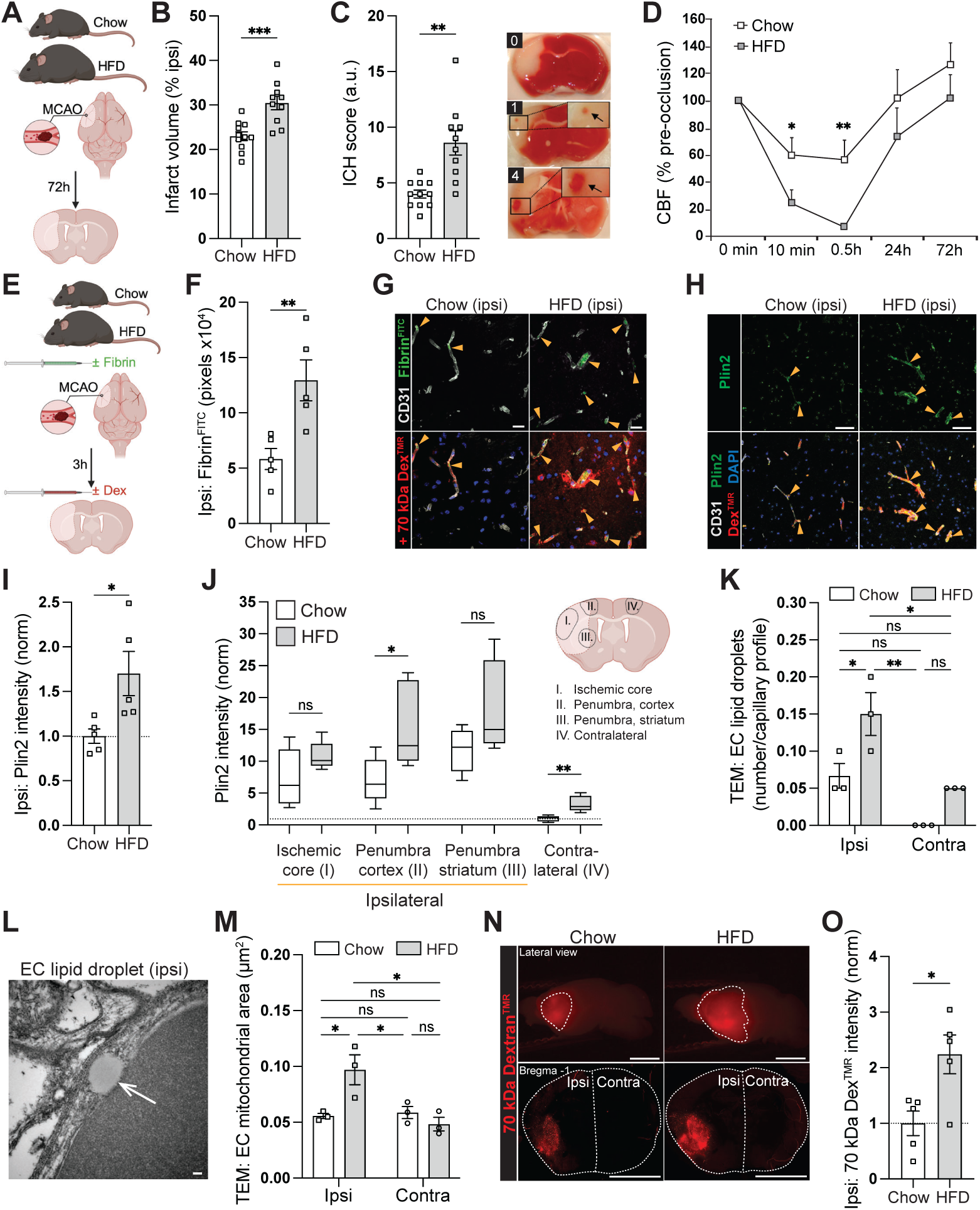
High-fat diet feeding confers a pro-thrombotic and reperfusion-resistant phenotype after acute ischemic stroke, characterized by vascular hyperpermeability and ectopic endothelial lipid droplet accumulation. **(A)** Chow and HFD mice subjected to MCAO (50 mg/kg Rose Bengal) and sacrificed 72 h later. **(B)** Infarct volumes determined after TTC staining. **(C)** Petechial hemorrhages counted and summed up to generate an ICH score. Representative score 0, 1 and 4 to the right. **(D)** CBF measured from the MCA territory in mice undergoing a milder MCAO protocol (12.5 mg/kg Rose Bengal). **(E)** Schematic representation of experimental outline for investigation of thrombosis and BBB function 3 h post MCAO (50 mg/kg Rose Bengal). **(F-G)** Microthrombi amount (F) and localization (G) in leaky blood vessels. **(H)** Cerebrovascular LDs in leaky blood vessels. **(I)** Overall Plin2 intensity quantified from ipsilateral hemisphere. **(J)** Plin2 quantified in three discrete ipsilateral regions: Cortical ischemic core, cortical-(dorsal) and striatal (medio-ventral) penumbra, and the corresponding dorsal cortical region in the contralateral hemisphere. **(K)** Quantification of endothelial LDs. **(L)** TEM image showing an endothelial LD (arrow). **(M)** EC mitochondria area. **(N)** Dex^TMR^ extravasation in whole-mount views (upper panels) and histological cross-sections (lower panels). **(O)** Dex^TMR^ intensity quantified in histological cross-sections. Scale bars, 50 μm (G-H); 500 μm (N); 1 nm (L). Data presented as mean ± s.e.m of n=10-11 mice/group (B-C), n=5 mice/group (D, F, I, O) and n=3 mice/group (K, M). Statistical evaluation using unpaired *t*-test (B, D, F, J, O), unpaired *t*-test with Welch’s correction (C, I) or two-way ANOVA followed by Holm-Šídák’s multiple comparisons test (K, M). \**P*<0.05, \*\**P*<0.001, \*\*\**P*<0.001. See also Figure S3.

Fibrin-congested vessels exhibited increased vasogenic permeability (**Figure 3G**). Intriguingly, oxygen deprivation has been shown to induce FA uptake and neutral lipid storage in cancer cells.^38^ As our *in vitro* data suggested that LD accumulation impairs EC integrity, we investigated whether ischemic stroke promotes LD synthesis. At 3 h post-MCAO, prominent LD accumulation was observed in leaky cerebral vessels (**Figure 3H**). HFD feeding increased overall Plin2 expression in the ischemic as well as non-ischemic hemisphere (**Figure 3I and 3J**), collectively suggesting that both hypoxia and circulating NEFAs may drive cerebrovascular LD formation after stroke. Furthermore, endothelial LD accumulation correlated with mitochondrial swelling (**Figures 3K, 3L and 3M**) indicating oxidative stress. Vascular hyperpermeability is known to exacerbate brain injury after stroke.^30,39^ At 3 h post-MCAO, when IgG and exogenous tracers readily penetrate the BBB^40^, we observed a more compromised BBB integrity in HFD mice (**Figures 3N and 3O; Figures S3I, S3J, S3K and S3L**). Hence, ectopic LD accumulation in the ischemic cerebrovasculature after MCAO, and the significant increase in response to HFD, intriguingly connects an augmented FA flux and increased plasma NEFA and TG levels with EC stress and deterioration of BBB integrity and ICH after stroke.

A modest peri-infarct *Vegfb* upregulation at 3 h post-MCAO progressed after 24 hours in HFD mice, indicative of a locally induced environmental cue regulating VEGF-B expression specifically in the penumbra (**Figure S3M**). No change in overall *Vegfb* or *Flt1* (VEGFR1) expression was recorded when analyzing RNA purified from whole hemispheres at 3 h post-MCAO (**Figures S3N, S3O and S3P**). *Nrp1* (neuropilin-1) expression was modestly upregulated in ischemic hemisphere in chow but not HFD mice (**Figure S3Q**), in agreement with previous studies.^41^ These data suggest a limited impact on local VEGF-B signaling after stroke. Taken together, the observed increase in infarct volume and ICH in HFD mice are likely explained by higher plasma lipids predisposing to pro-thrombogenic properties and augmented cerebrovascular ectopic LD accumulation, respectively.

### Pre-treatment with neutralizing anti-VEGF-B antibodies improves outcome after ischemic stroke by improving reperfusion and blood-brain barrier integrity, correlating with decreased cerebrovascular lipid droplet accumulation

Short-term (one week) αVEGF-B pre-treatment markedly reduced infarct volume and ICH events in HFD mice (**Figures 4A, 4B and 4C**). Pre-treatment with a humanized version of the αVEGF-B antibody developed for clinical use reproduced these beneficial effects (**Figures S4A, S4B and S4C**). A beneficial, but smaller, effect was observed also in chow mice (**Figures 4A, 4B and 4C**). Laser speckle contrast imaging at 3 h post-MCAO showed reduced penumbral blood perfusion in HFD *vs.* chow mice, and improved penumbra perfusion in αVEGF-B pre-treated HFD mice (**Figures 4D, 4E and 4F**). Preservation of BBB integrity after αVEGF-B pre-treatment was observed in both chow and HFD mice (**Figures 4G and 4H; Figures S4D, S4E, S4F, S4G, S4H and S4I**). Together, these data demonstrate improved peri-infarct perfusion and an early functional restoration of BBB integrity in response to αVEGFB pre-treatment, likely contributing to the later beneficial effect on infarct size and ICH.

**Figure 4.**
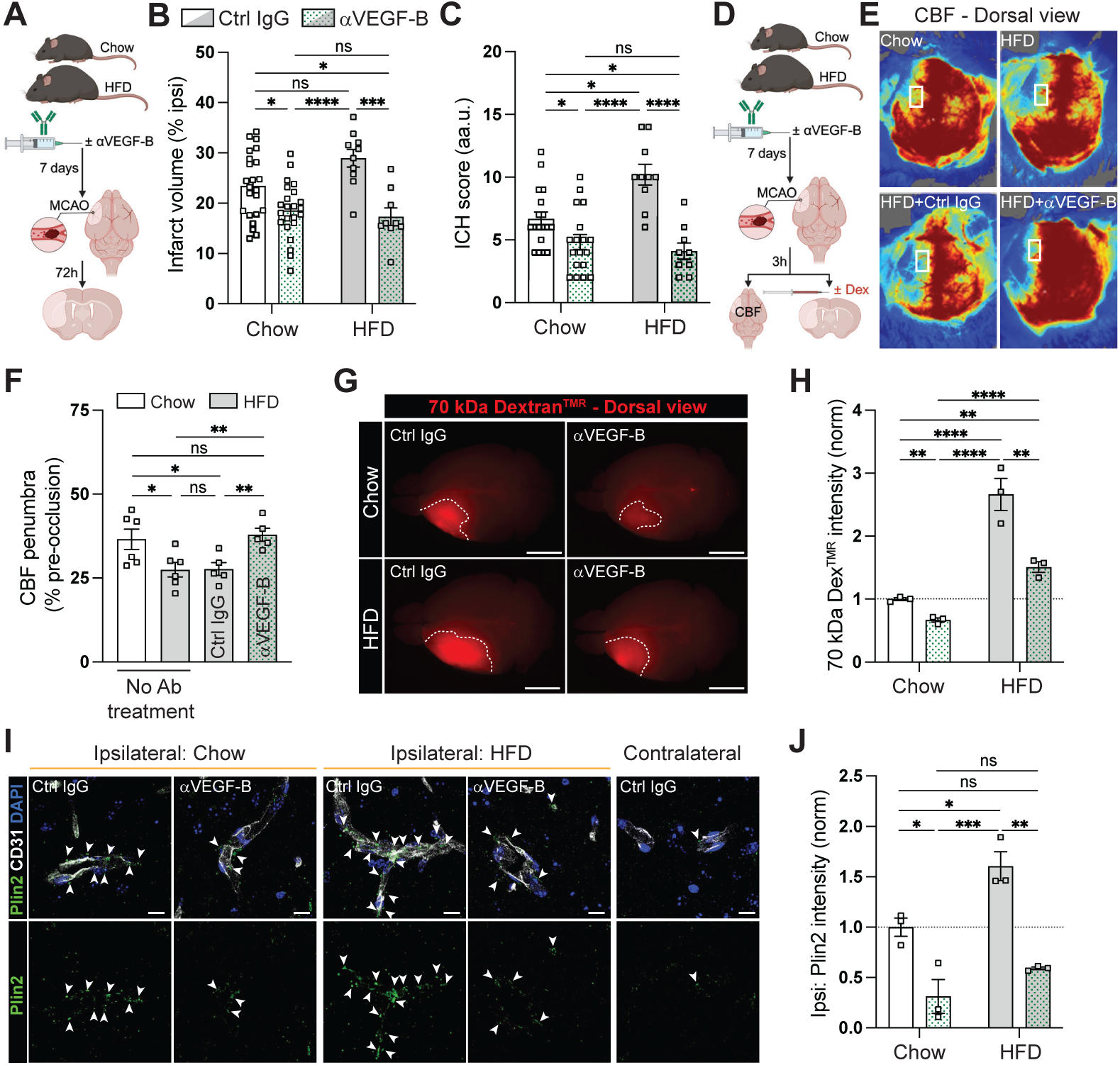
Pre-treatment with neutralizing anti-VEGF-B antibodies improves outcome after ischemic stroke by improving reperfusion and blood-brain barrier integrity, correlating with decreased cerebrovascular lipid droplet accumulation. **(A)** Chow or HFD mice pre-treated for 1 week with αVEGF-B or isotype control IgG antibodies before MCAO (50 mg/kg Rose Bengal) and sacrificed 72 h later. **(B)** Infarct volumes. **(C)** ICH scores. **(D)** Schematic representation of experimental outline in mice sacrificed 3 h post MCAO. **(E-F)** Evaluation of CBF with laser speckle contrast imaging (E). Boxed areas in the penumbra region utilized for quantification (F). (**G-H)** Dex^TMR^ extravasation in whole-mount views (G) and quantification (H). **(I-J)** Cerebrovascular Plin2+ LDs. Contralateral hemisphere shown for reference (I). Scale bars, 5 mm (G);10 μm (I). Data are presented as mean ± s.e.m. based on n=23+9-10 (chow *vs.* HFD) mice/group (B-C), n=5-6 mice/group (F) and n=3 mice/group (H, J). Statistical evaluation using two-way ANOVA followed by Tukeýs multiple comparisons test or one-way ANOVA followed by Fisheŕs LSD multiple comparisons test (F). \**P*<0.05, \*\**P*<0.01, \*\*\**P*<0.001, \*\*\*\**P*<0.0001. See also Figures S4 and S5.

Fewer cerebrovascular LDs and overall reduced Plin2 intensity in the ischemic hemisphere at 3 h post-MCAO was observed in αVEGF-B pre-treated chow and HFD mice (**Figure 4I and 4J**). Interestingly, Plin2+ LDs seemed to translocate from ECs to abluminal perivascular cells over time (**Figures S5A and S5B**), and accumulation of LDs in parenchymal cells reminiscent of lipid-laden microglia^42^ were appearing 24 h post-MCAO (**Figure S5C**), together indicating that LDs are dynamically metabolized. Interestingly, the reduction in ectopic LD accumulation was sustained at 24 h post-MCAO (**Figure S5D**). Global *Vegfb* knock-out mice on HFD exhibit decreased plasma NEFA and TG levels^43^, consistent with long-term^21,23^ and short-term (**Figures 2B and 2C**) αVEGF-B antibody pre-treatment. ICH was significantly decreased in *Vegfb*^-/-^ HFD mice, whereas infarct volume did not differ between *Vegfb*^-/-^ and littermate *Vegfb*^+/+^ controls (**Figures S5E, S5F and S5G**). The lack of a reduction in infarct size may be due to an opposing neuroprotective activity of VEGF-B within the brain parenchyma that is not inhibited by circulating αVEGF-B antibodies but is evident with congenital *Vegfb* deficiency. Based on these data we speculate that the primary beneficial effects of αVEGFB pre-treatment are a consequence of its lowering of plasma lipids leading to reduced cerebrovascular LD formation which would preserve BBB integrity and enhance glucose delivery.

### Therapeutic treatment with neutralizing anti-VEGF-B antibodies improves efficacy and safety of 5 h delayed intravenous thrombolysis in HFD mice

Both metabolic syndrome and delayed (>4.5 h post-stroke) reperfusion with rtPA correlates with hemorrhagic complications and worse outcome after ischemic stroke, limiting thrombolysis use in clinical stroke management.^2–4,26^ To evaluate the therapeutic potential of VEGF-B neutralization in a clinically relevant setting, antibodies were administered 1 h after confirmed MCA occlusion in HFD mice, followed by intravenous thrombolysis with rtPA at 5 h post MCAO (**Figure 5A**). Remarkably, a single injection of the αVEGF-B antibody after establishment of brain ischemia significantly reduced infarct volume, ICH, and severe hemorrhagic events intermittently observed after delayed thrombolysis (**Figure 5B, 5C and 5D**). A strong correlation between infarct size and ICH was observed (**Figure 5E**), where control-treated mice showed the highest propensity of ICH, presumably due to aggravated BBB damage triggered by HFD in combination with delayed rtPA reperfusion. The risk of fatal sICH coupled to delayed thrombolysis was further supported by the decreased survival in control-treated mice (**Figure 5F**), collectively suggesting that therapeutic αVEGF-B treatment confers improved safety and efficacy of delayed thrombolysis in HFD mice.

**Figure 5.**
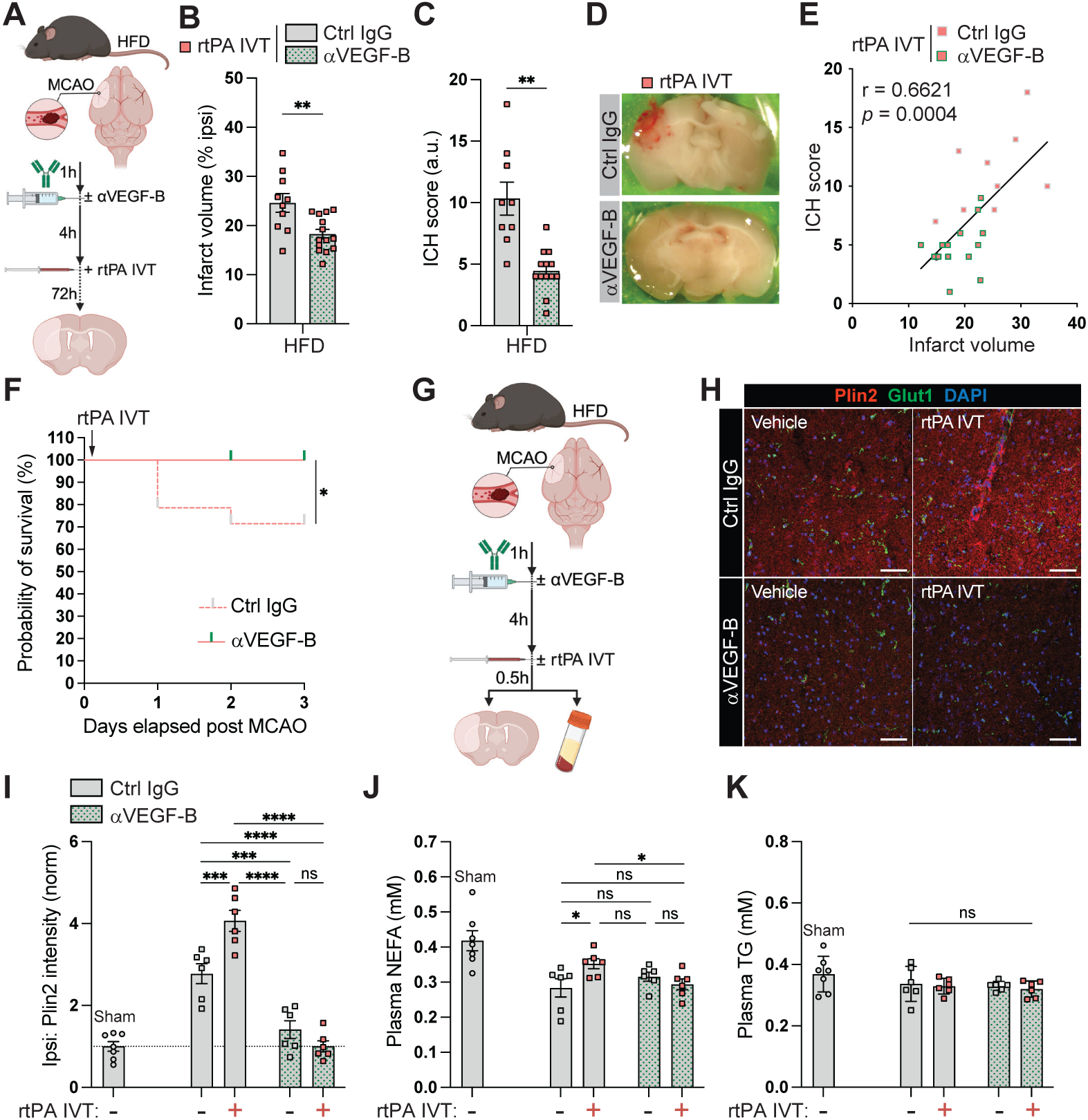
Therapeutic treatment with neutralizing anti-VEGF-B antibodies improves efficacy and safety of 5 h delayed intravenous thrombolysis in HFD mice. **(A)** HFD mice subjected to MCAO (50 mg/kg Rose Bengal) and 1 h later given one i.p. injection with αVEGF-B or isotype control antibodies, followed by intravenous thrombolysis (IVT) with rtPA (alteplase, 10 mg/kg) 5 h post MCAO. Experimental endpoint at 72 h post MCAO. **(B)** Infarct volumes. (**C)** ICH scores. **(D)** Hemorrhagic transformation in control, but not αVEGF-B treated mice. **(E)** Correlation between ICH events and infarct size. **(F)** Kaplan-Meier survival analysis. **(G)** Experimental endpoint at 5.5 h post MCAO including non-IVT control groups and sham mice as control for surgery and post-op procedures. **(H-I)** Plin2+ LD distribution (H) and quantification (I). **(J-K)** Plasma NEFAs (J) and plasma TGs (K). Scale bars, 50 μm. Data are presented as mean ± s.e.m. based on n=10+14 (Ctrl IgG *vs.* αVEGF-B) mice/group (B-C, E), n=14 mice/group (F) and n=6-7 mice/group (I-K). Statistical evaluation using unpaired *t*-test (B), unpaired *t*-test with Welch’s correction (C), linear regression with Pearson correlation (E), Log-rank (Mantel-Cox) test (F) and two-way ANOVA followed by Fisheŕs LSD test (I-K). \**P*<0.05, \*\**P*<0.01, \*\*\**P*<0.001, \*\*\*\**P*<0.0001. See also Figure S6.

Post-occlusion administration of the αVEGF-B antibody alone, omitting recanalization with rtPA, showed no effect on infarct volume or ICH (**Figures S6A, S6B and S6C**), suggesting that therapeutic αVEGF-B treatment specifically protects against thrombolysis-induced injury. Despite being ineffective on outcome measures at 72 h post-MCAO, LD formation was decreased and BBB integrity improved in the ischemic hemisphere 4 h post-αVEGF-B treatment (**Figures S6D, S6E, S6F, S6G, S6H, S6I and S6J**), validating the link between cerebrovascular LD accumulation and vascular hyperpermeability. Absent effect on circulating lipids (**Figures S6K, S6L, S6M and S6N**), however, indicated a local, rather than systemic, effect on brain LD synthesis. Hence, without concomitant restoration of blood flow, post-occlusion administration of αVEGF-B antibodies fail to improve outcome despite a diminishing effect on LD accumulation and preservation of BBB integrity acutely after MCAO.

The selective efficacy when combined with thrombolysis pointed towards a VEGF-B-dependent adverse activity exerted by rtPA. Intriguingly, LD formation in the ischemic hemisphere was exacerbated 30 min after thrombolysis (**Figures 5G, 5H and 5I**), coinciding with increased plasma NEFAs, but not TGs (**Figures 5J and 5K**), indicating that intravenous rtPA administration activates a systemic FA flux, analogous to the mechanism of action of VEGF-B.^23^ In support of a VEGF-B-dependent effect of rtPA, therapeutic αVEGF-B treatment abolished rtPA-induced LD accumulation in the ischemic hemisphere (**Figures 5H and 5I**), as well as the rise in plasma NEFAs **(Figure 5J).** Circulating NEFAs are readily taken up by the liver after WAT lipolysis.^23^ In accordance, Plin2+ LDs increased in liver after thrombolysis (**Figures S7A, S7B and S7C**). Collectively, these data demonstrate that limiting VEGF-B signaling counteracts rtPA-induced plasma NEFA rise and LD buildup in brain.

### Intravenous thrombolysis with rtPA activates visceral adipose tissue lipolysis

The acute and selective rise in plasma NEFAs, but not TGs, after thrombolysis with rtPA indicated an activation of WAT lipolysis. Lipolysis rate is regulated by phosphorylation of the LD-associated protein hormone-sensitive lipase (HSL).^44^ Indeed, phosphorylation of HSL increased in visceral WAT depots 30 min after rtPA thrombolysis (5.5 h post MCAO) compared to mice receiving vehicle control (**Figures 6A, 6B, 6C and 6D**). Interestingly, induction of MCAO itself promoted HSL activation, indicating an actively ongoing lipolysis acutely after stroke, presumably as part of a general sympathetic stress response.^45,46^ αVEGF-B co-treatment completely abolished rtPA-induced HSL activation and breakdown of stored TGs in visceral WAT (**Figures 6A, 6B, 6C and 6D; Figure S7D**), supporting a VEGF-B-dependent effect by rtPA specifically in visceral WAT.

**Figure 6.**
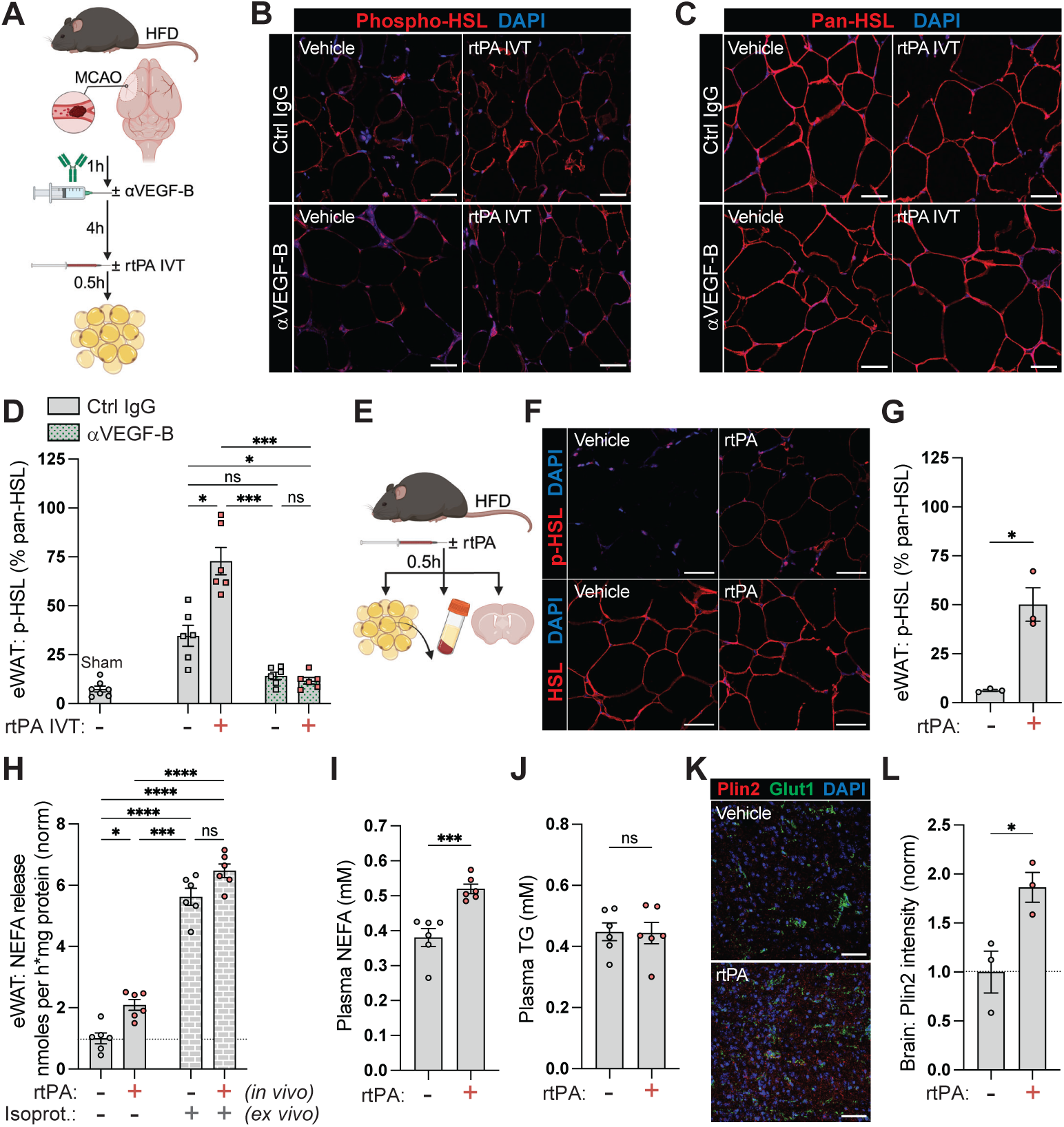
Intravenous thrombolysis with rtPA activates visceral adipose tissue lipolysis. **(A)** HFD mice subjected to MCAO (50 mg/kg Rose Bengal) and 1 h later given one i.p. injection with αVEGF-B or isotype control antibodies, followed by IVT with rtPA (alteplase, 10 mg/kg) or vehicle as control, at 5 h post MCAO. Experimental endpoint at 5.5 h post MCAO. Sham mice included as control for surgery and post-op procedures. **(B-C)** Activated (phosphorylated, B) and total (pan, C) hormone-sensitive lipase (HSL) in epididymal WAT (eWAT). **(D)** Phosphorylated/total HSL ratios. **(E)** Intravenous rtPA (alteplase, 10 mg/kg) or vehicle control injection of naïve HFD mice. Experimental endpoint 30 min post injection. **(F-G)** Phosphorylated/total HSL expression (F) and ratios (G) in eWAT. **(H)** *Ex vivo* lipolysis in eWAT (basal and isoproterenol stimulated). **(I-J)** Plasma NEFAs (I) and TGs (J). **(K-L)** Plin2+ LD distribution (K) and quantification (L) in brain. Scale bars, 50 μm. Data presented as mean ± s.e.m. based on n=6-7 mice/group (D, H-J) and n=3 mice/group (G, L). Statistical evaluation using Brown-Forsythe and Welch ANOVA followed by Dunnett’s T3 multiple comparisons test (D), two-way ANOVA followed by Tukeýs multiple comparisons test (H), unpaired *t*-test (I-J, L), or unpaired *t*-test with Welch’s correction (G). \**P*<0.05, \*\*\**P*<0.001, \*\*\*\**P*<0.0001. See also Figure S7.

Notably, intravenous rtPA administration activated HSL and decreased adipocyte cell size in visceral WAT independent on MCAO (**Figures 6E, 6F and 6G; Figures S7E and S7F**). To provide functional evidence for a rtPA-induced lipolytic effect, NEFA release was quantified from WAT depots collected 30 min after intravenous rtPA administration in naïve HFD mice. rtPA increased the rate of NEFA released from epididymal (**Figure 6H**) and perirenal (**Figure S7G**) WAT (both visceral), but not subcutaneous WAT (**Figure S7H**), in line with an effect primarily on metabolically active WAT.^47^ *Ex vivo* stimulation with isoproterenol, a nor-epinephrine (NE) mimic, revealed no additive or synergistic effect over rtPA, pointing towards a mechanism of action within the sympathetic pathway. Increased lipolysis correlated with a selective rise in plasma NEFAs (**Figures 6I and 6J**) and increased LD formation in brain (**Figures 6K and 6L**) and liver (**Figures S7I and S7J**), collectively demonstrating that intravenous rtPA administration, even in the absence of cerebral ischemia, promotes visceral WAT lipolysis, leading to a surge of circulating NEFAs and LD buildup in brain and liver.

### Blocking VEGF-B signaling in adipose tissue is sufficient to improve stroke outcome and prevent rtPA-induced lipolysis

Plasma NEFA concentration is dependent on the rate of WAT lipolysis and tissue uptake, respectively. Both HFD feeding and intravenous rtPA administration increased the circulating NEFA pool, correlating with poor outcome after MCAO. As systemic αVEGF-B treatment prevented both HFD- and rtPA-induced plasma NEFA increase, we next attempted to dissect whether the improved outcome after MCAO was attributable to the anti-lipolytic effect in WAT, or if local VEGF-B signaling in brain plays a central role in stroke outcome. Genetically engineered mice with reduced VEGF-B expression specifically in adipose tissue (Adipoq^Cre/+^;*Vegfb*^fl/+^) have lower plasma NEFAs compared to littermate controls, mechanistically linked to a reduced lipolytic rate and NEFA transport to blood.^23^ Moreover, WAT insulin sensitivity is markedly improved in Adipoq^Cre/+^;*Vegfb*^fl/+^, compared to control, HFD mice.^23^ Reducing VEGF-B signaling selectively in adipose tissue phenocopied the effect on stroke outcome after systemic VEGF-B neutralization (**Figures 7A, 7B and 7C**), signifying that targeting VEGF-B signaling explicitly in WAT, with direct impact on circulating plasma NEFAs, is sufficient to improve outcome after acute ischemic stroke.

**Figure 7.**
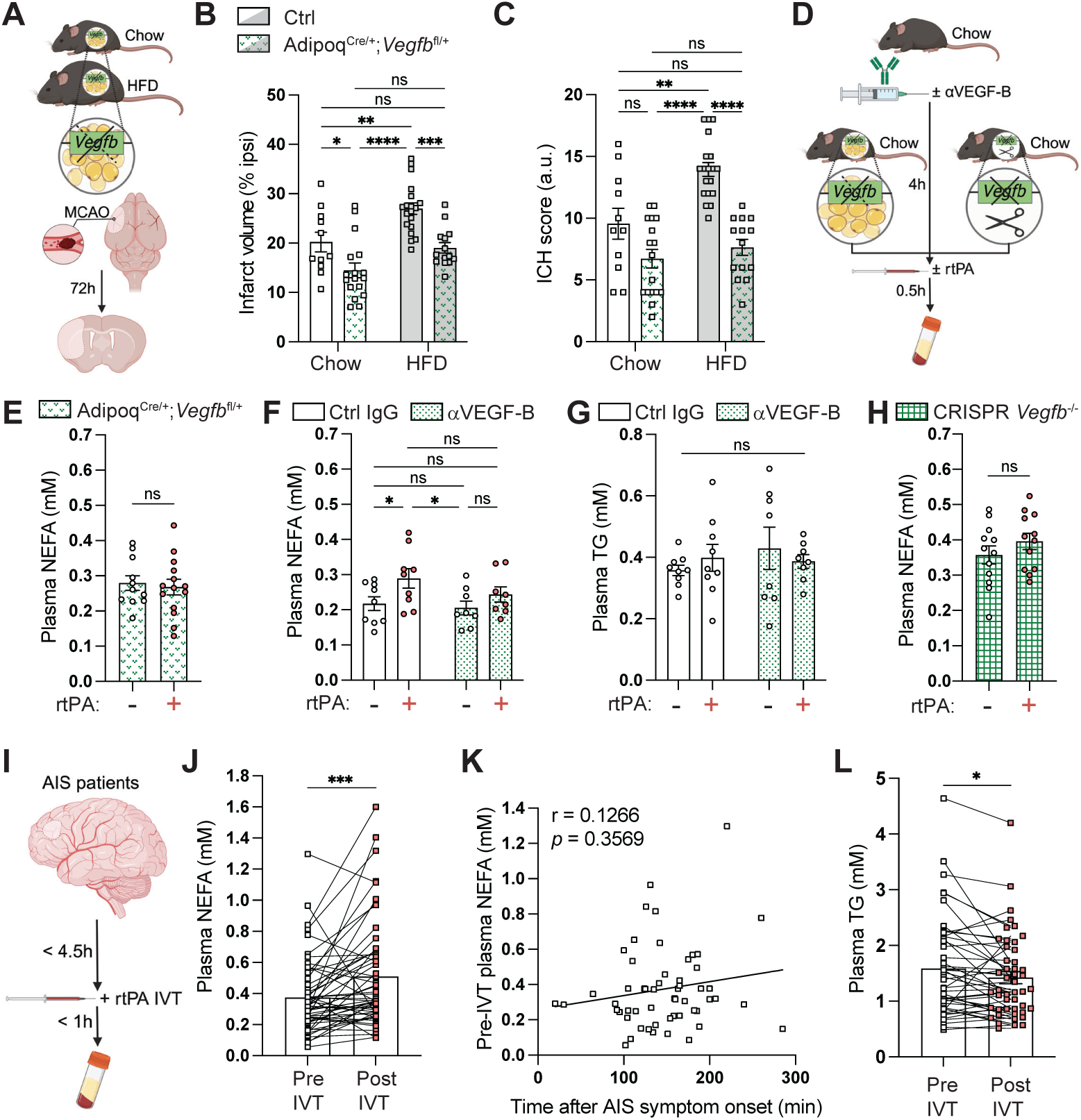
Blocking VEGF-B signaling in adipose tissue is sufficient to improve stroke outcome and prevent rtPA-induced lipolysis. **(A)** Chow or HFD-fed WAT-specific *Vegfb* haplo-insufficient mice: Adipoq^Cre/+^;*Vegfb*^fl/+^ and sibling controls subjected to MCAO (50 mg/kg Rose Bengal) and sacrificed 72 h later. **(B)** Infarct volumes. **(C)** ICH scores. **(D)** Intravenous rtPA (alteplase, 10 mg/kg), or vehicle as control, injection of naïve Adipoq^Cre/+^;*Vegfb*^fl/+^, congenital *Vegfb*^-/-^, or αVEGF-B *vs.* ctrl IgG pre-treated (4 h before) mice on chow diet. Experimental endpoint 30 min post rtPA injection. **(E)** Plasma NEFAs in naïve Adipoq^Cre/+^;*Vegfb*^fl/+^ mice. **(F-G)** Plasma NEFAs (F) and plasma TGs (G) after systemic pre-treatment with neutralizing αVEGF-B antibodies. (**H)** Plasma NEFAs in *Vegfb*^-/-^ mice. **(I)** Acute ischemic stroke patients eligible for intravenous thrombolysis (IVT) enrolled within 4.5 h post stroke symptom onset. Paired blood samples taken before thrombolysis, and within 1 h after completion of rtPA (alteplase) infusion. **(J)** Plasma NEFAs. **(K)** Correlation between duration of acute ischemic stroke symptoms and plasma NEFAs. **(L)** Plasma TGs. Data presented as mean ± s.e.m. based on n=11+17 (chow); 19+14 (HFD) (Ctrl *vs.* Adipoq^Cre/+^;*Vegfb*^fl/+^) (B-C), n=11+14 mice/group (E), n=8-9 mice/group (F-G), n=12 mice/group (H) and n=55 acute ischemic stroke patients (J-L). Statistical evaluation using two-way ANOVA followed by Tukeýs multiple comparisons test (B-C), unpaired *t*-test (E, H), two-way ANOVA followed by Fisheŕs LSD test (F), Brown-Forsythe and Welch ANOVA followed by Dunnett’s T3 multiple comparisons test (G), linear regression with Wilcoxon matched-pairs signed rank test (J, L) or Pearson correlation (K). \**P*<0.05, \*\**P*<0.01, \*\*\**P*<0.001, \*\*\*\**P*<0.0001. See also Figure S7.

An abolished effect on plasma NEFAs 30 min after intravenous rtPA administration in chow fed Adipoq^Cre/+^;*Vegfb*^fl/+^ mice (**Figure 7E**) confirmed and advanced the notion that rtPA-induced lipolysis is regulated by VEGF-B in WAT. Systemic treatment with αVEGF-B antibodies 4 h before rtPA administration also prevented plasma NEFA rise (**Figure 7F**), confirming that circulating antibodies readily access WAT and inhibits rtPA-induced lipolysis. Plasma TGs remained unaffected (**Figure 7G**), as previously observed in HFD mice. Likewise, *Vegfb*^-/-^ mice were refractory to rtPA-induced lipolysis (**Figure 7H**), together illustrating the fundamental reliance on active VEGF-B signaling in WAT to exert rtPA-induced lipolysis.

### A thrombolysis-induced rise in plasma fatty acids is observed in ischemic stroke patients treated with rtPA

Plasma TGs are regularly monitored and used as a biomarker for stroke outcome in the clinic.^48,49^ Circulating NEFAs are however scarcely assessed. To examine the translational significance of rtPA-induced lipolysis, plasma parameters were evaluated in patients receiving rtPA (alteplase) thrombolysis. Matched plasma samples from 55 acute ischemic stroke patients, collected immediately before and <1 h after completion of thrombolysis (**Figure 7I**), revealed a net NEFA increase in response to rtPA treatment (**Figure 7J**). Pre-thrombolysis plasma NEFAs did not significantly increase with time after stroke symptom onset (**Figure 7K**), indicating a true rtPA-induced effect. The simultaneous decrease in plasma TGs (**Figure 7L**) further supported the concept of an acute pro-lipolytic response after thrombolytic therapy.

## Discussion

Intravenous thrombolysis and endovascular thrombectomy are two evidence-based treatments for acute ischemic stroke.^24^ Endogenous tPA secreted from ECs activates plasmin-dependent fibrinolysis, providing the underlying mechanistic rationale for re-canalization with rtPA. Thrombolytic therapy with rtPA provides an overall long-term benefit of reduced disability, with a net favorable benefit/risk ratio.^25,26,50^ However, it also carries an early risk of ICH and hemorrhagic transformation that is increased with delayed rtPA treatment (>4.5 h after stroke symptom onset). This risk is further exacerbated by metabolic syndrome features, such as obesity, hyperglycemia, and dyslipidemia.^2–4,8,10^ The underlying mechanisms driving these increased risks are not fully understood. Thus, if the incidence or magnitude of ICH could be diminished, the benefit/risk ratio after thrombolysis could be significantly improved. In the current paper we describe novel mechanistic insights into the systemic influences of rtPA on ICH and stroke outcome, involving an unexpected acute lipolytic activity mediated by rtPA. Of potential translational significance, we show that adjuvant treatment with a neutralizing αVEGF-B antibody blocked thrombolysis-induced lipolysis and extended the time window for safe and efficacious re-canalization with rtPA.

Glucose constitutes the essential and predominant energy substrate for the brain and glucose delivery relies on GLUT1-mediated uptake in BBB ECs.^51,52^ Altered brain glucose utilization in hyperglycemia and metabolic syndrome has been implied, albeit no consensus conclusion has so far been reached regarding the particular features.^51,53–56^ We specifically found that FA exposure impaired EC glucose uptake, lending a possible explanation to the observed reduced glycolytic rate in FA-treated ECs.^57^ ECs predominantly utilize glucose for their own metabolic needs^33,52,58^ and ECs exposed to long-chain FAs instead upregulated FA uptake. That FA flux regulates systemic glucose metabolism is evident for many factors controlling endothelial long-chain FA uptake, including VEGF-B^16,21,43^, 3-hydroxyisobutyrate^59^, lactate^60^, angiopoietin-2^61^, apelin^62,63^, delta-like canonical notch ligand 4 (Dll4)/notch^64^ and peroxisome proliferator-activated receptor-ψ (PPARψ).^65^ Upregulation of VEGF-B expression induces FA uptake from blood via paracrine VEGFR1/neuropilin-1 signaling in ECs and subsequent buildup of LDs in *e.g.* cardiac tissue, correlating with a corresponding decrease in glucose uptake rate.^16,33^ Intriguingly, WAT ECs are involved in bidirectional transport, where FAs are transported in the reverse direction after intracellular lipolysis, contributing to plasma NEFA rise. Of importance for the mechanistic studies included in this paper, reducing VEGF-B signaling selectively in WAT is sufficient to prevent lipolysis-induced plasma NEFA rise and these mice also exhibit overall decreased circulating NEFAs.^23^ Global VEGF-B inhibition using neutralizing antibodies or whole-body gene knock-out strategy, however, will in addition decrease NEFA uptake from blood, resulting in net lowering of plasma NEFAs predominantly in conditions characterized by augmented lipolysis, for example fasting and HFD.^21,23,43^ HFD-induced insulin resistance promotes hyperglycemia and dyslipidemia, characterized by elevated plasma NEFAs derived from augmented WAT lipolysis.^23^ Intriguingly, we observed decreased cerebral glucose uptake/utilization in HFD mice, in line with a previous report showing correlation between increased insulin resistance and decreased brain glucose uptake/utilization.^66^ We however did not find consistent changes in *Slc2a1* (Glut1) expression *in vivo*, suggesting an alternative regulation of brain glucose uptake. Of further relevance to our study, augmented long-chain FA uptake in brain has been reported in metabolic syndrome patients.^67,68^ αVEGF-B pre-treatment reduced plasma NEFAs, correlating with improved glucose uptake and less dense LD distribution in brain, signifying a connection between increased FA flux and reduced brain glucose uptake, in line with the established concept of impaired glucose uptake correlating with ectopic lipid deposition.^69^

Increased infarct volume in HFD mice has previously been observed with transient MCAO models, suggestively attributed to an altered inflammatory response.^36,37^ We observed increased thrombosis and refractory endogenous fibrinolysis in our photothrombotic MCAO model. Long-chain FAs promote thrombogenesis^70^ and we therefore postulate that elevated plasma NEFAs contributed to enhanced secondary thrombotic events in HFD mice which in turn resulted in increased infarct volume.^71^ In line with this hypothesis, the greater effect size on infarct volume reduction after αVEGF-B pre-treatment in HFD, compared to chow mice, could be mechanistically linked to plasma NEFA lowering. That stroke outcome is influenced by circulating NEFAs was further supported by that WAT-specific *Vegfb* haploinsufficiency phenocopied the αVEGF-B antibody effect, notably also without affecting blood glucose levels.^23^

Increased plasma NEFAs correlate with ectopic LD deposition ^21,23,43,72^, but FA flux and LD buildup after ischemic stroke has not been explored in detail previously. Within 3 hours post-MCAO, LDs accumulated in medium- to large caliber vessels in ischemic core and penumbra. Cerebrovascular LDs have previously been observed 24 h after transient MCAO, correlating with a robust inflammatory gene signature and upregulation of the FA importer CD36 and LD marker Plin2 in isolated brain ECs.^42^ Our data extend this observation by determining that ectopic cerebrovascular LD accumulation occur earlier than previously reported and moreover, proportionally to circulating NEFA levels. Interestingly, lactate has been shown to increase long-chain FA uptake and LD buildup in ECs.^60^ Activation of anaerobic glycolysis and enhanced lactate production in the ischemic hemisphere could thus conceivably drive endothelial FA uptake and contribute to the distinct accumulation of LDs specifically in the ischemic hemisphere. *Vegfb* upregulation in the penumbra, possibly as a response to glucose starvation downstream PGC1α^43^, may also have contributed. Our data however indicate that high FA content in blood can drive the shift from glucose to FA uptake in brain independently on local VEGF-B signaling. Notably, FA overload, overriding endothelial LD storage capacity and disposal through beta-oxidation, is coupled to lipotoxicity.^54,57,58^ LD-loaded ECs display ER stress, augmented proinflammatory signaling and decreased nitric oxide synthesis, correlating with endothelial activation and defective vascular relaxation.^73,74^ Therefore, αVEGF-B treatment could apart from enhancing glucose delivery also improve vasodilatory capacity, a plausible explanation to the improved blood perfusion.

Increased endothelial FA uptake and LD formation correlated with augmented trans- and paracellular permeability *in vitro*. In support of our findings, exposure to long-chain FAs, but interestingly not high glucose, has been shown to compromise brain EC integrity.^75^ Features of the metabolic syndrome, mainly attributed to hyperglycemia, is connected to vasogenic edema and increased propensity of ICH after stroke.^2,32,36,53,55,76^ αVEGF-B pre-treatment improved BBB integrity which we propose is connected to lowering of plasma NEFAs and decreased FA uptake and LD accumulation. The early permeability peak (hours post-MCAO) is followed by a more severe second wave (days post-MCAO), where basal lamina breakdown and glia barrier perturbation may lead to ICH and hemorrhagic transformation.^32^ In support of that early vulnerability of the BBB, coupled to increased LD formation, is predisposing for ICH later on was that αVEGF-B pre-treatment ameliorated both early vascular hyperpermeability and ICH. Global *Vegfb*^-/-^ mice are conceivably not particularly useful to study neuronal survival after stroke due to a congenital defect in neurogenesis.^77^ *Vegfb*^-/-^ HFD mice however reproduced amelioration of ICH after MCAO, supporting the notion that plasma lipid lowering confers BBB protection. The smaller, yet significant, ameliorating effect of αVEGF-B pre-treatment in chow mice, despite unaffected plasma NEFAs, might be connected to inhibition of cerebrovascular VEGF-B signaling, conferring reduced FA uptake from blood. In line with this assumption, overall effect size was smaller in mice harboring WAT-specific *Vegfb* haplodeficiency, compared to antibody pre-treated mice.

Intravenous thrombolysis with rtPA increases the risk of sICH and hemorrhagic transformation after stroke.^27^ In animal models, hemorrhagic complications associated with thrombolysis has been shown to depend on increased neutrophil recruitment and degranulation^78^, as well as matrix metalloproteinase 9 (MMP9) activity^79^, attributed to local effects of rtPA interacting with the damaged cerebrovasculature after recanalization. Furthermore, we have earlier demonstrated glial barrier perturbation downstream local activation of platelet-derived growth factor -CC (PDGF-CC) by tPA, connected to ICH after MCAO.^28–30^ Limiting FA flux as an adjuvant co-treatment to rtPA thrombolysis has however previously not been assessed. Five-hour delayed thrombolysis in HFD mice correlated with reduced survival. The underlying cause of this increased mortality need to be determined, but majority of mice died 24 h post MCAO, most likely due to hemorrhagic complications. In comparison, 100% of the mice receiving therapeutic αVEGF-B antibodies survived thrombolytic intervention until the experimental endpoint and exhibited ameliorated ICH and reduced infarct size. To predict translational outcome even better, evaluation of neurological deterioration in the acute phase as well as long-term functional efficacy of adjuvant combination treatment with thrombolysis is warranted.

Apart from expression in ECs, tPA is co-expressed and co-released with NE in sympathetic nerves.^80^ tPA inhibition/deficiency attenuates NE release and sympathetic responses^81^, whereas tPA/rtPA treatment induces NE release, collectively proposing a potentiating effect on NE activity.^80–82^ A major novel finding in this study was the discovery of an acute pro-lipolytic activity of intravenously administered rtPA in visceral WAT; via activation of HSL, a main regulated and rate limiting enzyme involved in TG hydrolysis from intracellular LDs.^44^ No additive effect was observed on top of NE stimulation, suggesting that rtPA either releases a break on basal lipolysis or stimulates lipolysis within the NE pathway. It remains yet to dissect whether this effect is dependent on tPÁs proteolytic activity and if rtPA acts within the VEGF-B signaling pathway. That the pro-lipolytic rtPA effect is reversible by limiting VEGF-B signaling specifically in WAT was however confidently concluded using WAT-specific *Vegfb* haploinsufficient mice. Therapeutic (post-MCAO) administration of the αVEGF-B antibody abolished thrombolysis-induced WAT lipolysis and plasma NEFA rise. Adjuvant post-treatment also prevented LD accumulation in the ischemic hemisphere, which associated with improved BBB integrity. Increased plasma NEFAs at 24 h post-stroke has been observed in a genetic obesity mouse model^83^ and our data demonstrate induced WAT lipolysis and signs of enhanced systemic FA flux already within hours after an ischemic insult to some extent also in the absence of thrombolysis. Increased liver LD synthesis after thrombolysis is presumably driven by sequestering of circulating NEFAs. A direct effect of MCAO or rtPA on *de novo* lipogenesis in liver can however not be excluded. NEFA uptake in liver is independent on local VEGF-B signaling^16,23^, possibly explaining the stronger LD-inhibiting effect in brain, compared to liver, after systemic VEGF-B inhibition. Build-up of transient or even permanent neutral lipid storage in liver after stroke has, to our knowledge, not been studied. Without recanalization with rtPA, αVEGF-B post-treatment did not ameliorate outcome measures at 72 h post-MCAO, and we speculate that this is due to neither thrombosis nor spontaneous reperfusion being affected by post-treatment alone. Our data thus point out that therapeutic treatment with αVEGF-B antibodies specifically ameliorate reperfusion injury after thrombolysis connected to increased WAT lipolysis and plasma NEFAs.

Acute ischemic stroke patients undergoing rtPA thrombolysis displayed increased plasma NEFA levels. The TyG (TG-glucose) index has emerged as a novel biomarker of insulin resistance and a high TyG index correlates with increased mortality, poor functional outcome, and lack of neurological improvement in stroke patients undergoing thrombolysis; however, no association to sICH was found.^48,49^ We advocate for an emphasis on also monitoring circulating NEFAs, a lipid species not regularly assessed in the clinic, as our data suggests that increased plasma NEFAs could predict ICH after thrombolysis. A larger clinical sample size is however needed to verify if plasma NEFAs correlate with incidence of sICH. In summary, an ideal adjunct treatment for rtPA should not only reduce ICH but also exert neuroprotective effects. Limiting lipolysis and FA flux with an αVEGF-B antibody at the time of thrombolysis decreased both infarct size and ICH and may constitute a future strategy to expand the therapeutic window and increase the number of patients eligible for rtPA treatment.

## Acknowledgements

This work was supported by grants from the Swedish Heart and Lung foundation and Diabetes Foundation (I.N., U.E.), Swedish Brain Foundation, FSG FANG FOUNDATION and Hållsten Research Foundation (I.N., L.F., U.E.), Swedish Governmental Agency for Innovation Systems (VINNOVA) and CSL Innovation Ltd. (L.F., U.E.), Swedish Stroke Foundation, Royal Swedish Academy of Sciences and Magnus Bergvalĺs foundation (I.N.), EMBO Long-Term Fellowship co-funded by Marie Curie Actions (C.M.), Lendület “Momentum” Grant of the Hungarian Academy of Sciences (Z.B.), National Institutes of Health (R01-HL055374, R01-AG074552) and American Heart Association 19TPA34880040 (D.A.L), Leducq Foundation and Novo Nordisk Foundation (U.E.), and Karolinska Institutet. The monoclonal anti-VEGF-B antibody is proprietary to and generously provided by CSL Limited, Parkville, Victoria, Australia.

We thank Stefan Craciun, Mary Montenegro, Alice Nilsson, Anna Olverling, Tofan Taeri and Sofia Wittgren for technical assistance, and Annelie Falkevall and Annika Mehlem for fruitful discussions.

## Author Contributions

I.N. and E.J.S. designed, carried out, analyzed and interpreted most animal and biochemical experiments. L.F. carried out animal experiments. B.H.S and C.M. designed, carried out and analyzed *in vitro* experiments. Z.B., H.H. and R.L.M. provided and analyzed human plasma. B.H.S, C.S., F.C.N., M.Z., L.M., A-L.E.L. and P.D.S. performed biochemical experiments or assisted with animal experiments. L.L. and E.S. carried out ^18^F-DG PET. R.F.K. performed and analyzed TEM data. D.A.L. and U.E. designed and directed research, interpreted data and allocated resources. I.N. wrote the paper, with assistance from all authors.

## Methods

### Primary human endothelial cell culture

Primary human umbilical vein endothelial cells (HUVECs; PromoCell, passage <7) were expanded on gelatin-coated (0.1%) tissue culture (TC)-treated dishes (BD Falcon) in Endothelial Cell Growth Medium 2 including growth factors and 2% fetal bovine serum (FBS) (EGM-2; PromoCell) and 1% Penicillin-Streptomycin (Invitrogen) in a humidified cell incubator at 37°C, 5% CO_2_. A human EC model of obesity and diabetes was established by growing HUVECs to confluency on gelatin-coated 24-well TC plates or glass chamber slides (4-well or 8-well; BD Falcon) in EGM-2 and thereafter changing to growth factor-depleted (GFD) media (Endothelial Cell Basal Medium, PromoCell, containing 2.5% FBS and 1% Penicillin-Streptomycin) supplemented with FA-free bovine serum albumin (BSA, Sigma-Aldrich) conjugated to long-chain FAs (0.1 mM; mix of 0.05 mM oleic acid and 0.05 mM palmitic acid, Larodan) and glucose (20 mM D-Glucose, Merck), alone or in combination, for 2 h, 24 h or 5 days with daily exchange of medium. Alternatively, GFD media was supplemented with FA-free BSA-conjugated oleic- and palmitic acid mix (0.05 mM; 0.025 mM each), 0.05 mM oleic acid or 0.5 mM oleic acid, for the indicated time periods. The BSA final concentration was 0.1% (or 1% when using 0.5 mM oleic acid). For VEGF-B and oleic acid co-treatment experiments, HUVECs were pre-conditioned for 24 h in EGM-2 supplemented with FA-free BSA-conjugated oleic acid (0.5 mM). Cells were thereafter washed and incubated for 2 h in GFD media supplemented with 0.5 mM oleic acid. Thereafter, growth factor-starved HUVECs were treated with VEGF-B_186_ (50 ng/ml; Sf21 derived, R&D Systems) or vehicle control for 2 h in GFD media supplemented or not with 0.5 mM oleic acid. The neutralizing capacity of the αVEGF-B monoclonal antibody was evaluated by stimulating confluent HUVECs pre-starved for 2 h in GFD media with a fixed amount of VEGF-B_186_ (50 ng/ml; R&D Systems) for 2 h in GFD media, titrated against increasing amounts (equimolar, 3- and 9- molar excess) of αVEGF-B antibody CSL346. At the end of the experiments, glucose- or FA uptake, fluorescent probe- or immunofluorescence, or barrier integrity was assessed.

Primary human brain microvascular endothelial cells (HBMECs; PromoCell, passage <5) were employed for VEGF-B stimulation experiments. HBMECs were expanded on gelatin-coated (0.1%) TC-treated dishes in Endothelial Cell Growth Medium MV2 including growth factors and 5% FBS (EGM-MV2; PromoCell) and 1% Penicillin-Streptomycin in a humidified cell incubator at 37°C, 5% CO_2_. Stimulations were performed in 24-well TC plates when cells grown in EGM-MV2 reached confluence. Prior to 2 h stimulation with VEGF-B protein or neutralizing αVEGF-B antibodies, the cells were starved from growth factors for 2 h in GFD media (Endothelial Cell Basal Medium MV2, PromoCell, containing 2.5% FBS and 1% Penicillin-Streptomycin). Recombinant mouse VEGF-B_167_ (100 ng/ml; *E. coli* derived), VEGF-B_186_ (100 ng/ml; Sf21 derived, both from R&D Systems), or 2 µg/ml αVEGF-B antibody was utilized. At the end of the experiments, glucose- or FA uptake was assessed.

### Glucose and fatty acid uptake in primary human endothelial cells

To evaluate glucose uptake capacity, confluent HUVECs or HBMECs grown in 24-well TC plates were washed in Krebs-Ringer solution without glucose (120 mM NaCl, 24 mM NaHCO_3_, 4.8 mM KCl, 1.2 mM MgCl_2_, 1.2 mM KH_2_PO_4_ and 5 mM HEPES, pH 7.4) for 10 min at 37°C, 5% CO_2_. The cells were thereafter incubated for 20 min at 37°C, 5% CO_2_ in GFD medium supplemented with 5 mM L-glucose and 1 mM 2-[N-(7-nitrobenz-2-oxa-1,3-diazol-4-yl)amino]-2-deoxy-D-glucose (2-NBDG; Invitrogen). Finally, the cells were washed 3x with PBS/5 mM D-glucose and fixed with 4% PFA/5 mM D-glucose. For evaluation of FA uptake, confluent HUVECs or HBMECs were incubated with PBS/0.1% FA-free BSA supplemented with 20 µM of a long-chain FA analogue: BODIPY^TM^-C_12_ 500/510-C_1_ (4,4-difluoro-5-methyl-4-bora-3a,4a-diaza-s-indacene-3-dodecanoic acid; Invitrogen) or BODIPY^TM^-C_12_ 558/568 (4,4-difluoro-5-(2-thienyl)-4-bora-3a,4a-diaza-*s*-indacene-3-dodecanoic acid; Invitrogen) for 5-10 min at 37°C, 5% CO_2_. Finally, the cells were washed 3x with PBS/0.1% FA-free BSA and fixed with 4% PFA. Cells were imaged immediately after finishing the glucose or FA uptake assay with a Zeiss Axio Observer Z1 microscope equipped with a 20x long distance Plan+Neofluar fluorescence objective (N/A=0.4 Ph2 Korr μ/1.5) and an AxioCam MRm fluorescence CCD camera. Each condition was performed in triplicates and 10-15 images per well were analyzed with ImageJ or AxioVision (Carl Zeiss) software by setting threshold and measuring the integrated density per frame or intensity of pixel^2^. Mean values were normalized to control. Pooled results for repeated experiments are shown.

### Fluorescent probes and immunofluorescence in primary human endothelial cells

To visualize mitochondria in HUVECs grown for 24 h on 0.1% gelatin-coated glass chamber slides (4- or 8-well; BD Falcon) in FA-supplemented or control media, MitoTracker^TM^ Red CMXRos (0.25 mM final concentration; Invitrogen) was added to the media for 15 min before experiment endpoint. Cells were washed once in PBS and fixed for 10 min on ice with 4% PFA prior to mounting in Prolong gold antifade reagent with DAPI (Invitrogen). Representative maximum intensity projections (2D renderings) of 4 μm confocal z-stacks acquired with a confocal laser scanning microscope (LSM 700, Zeiss) equipped with a 63x Oil (N/A = 1.4) objective and the ZEN 2009 software (Carl Zeiss Microimaging GmbH) is presented.

For evaluation of transcellular permeability in HUVECs grown for 24 h on 0.1% gelatin-coated glass chamber slides (BD Falcon) in FA-supplemented or control media, 1.25 mg/ml tetramethyl-rhodamine (TMR)-conjugated 70 kDa dextran (D1818, Molecular Probes) was added to the media 15 min before experiment endpoint. TMR-conjugated 70 kDa dextran (Dex^TMR^) is taken up by macro-pinocytosis and increased amount of dextran-filled transcytotic vesicles represent enhanced macromolecular permeability. Cells were washed once in PBS and fixed for 10 min on ice with 4% PFA prior to mounting in Prolong gold antifade reagent (Invitrogen). Each condition was performed in quadruplicates and five maximum intensity projections (2D renderings) of 4 μm confocal z-stacks per well were acquired with a confocal laser scanning microscope (LSM 700, Zeiss) equipped with 20x (N/A=0.8) objective and the ZEN 2009 software (Carl Zeiss Microimaging GmbH) using the same settings (within the respective experiment). For quantification of Dex^TMR^ signal using integrated density, the number of pixels above a set threshold was determined using the ImageJ software. Mean values were normalized to control and pooled results for repeated experiments are shown.

For evaluation of intracellular neutral lipid content, HUVECs grown for 24 h on 0.1% gelatin-coated 24-well TC plates in FA-supplemented or control media were washed with PBS and fixed on ice for 10 min in 4% PFA. Neutral lipids were stained using 0.1 mg/ml BODIPY^TM^ 493/503 (4,4-Difluoro-1,3,5,7,8-Pentamethyl-4-Bora-3a,4a-Diaza-*s*-Indacene; Invitrogen) and 0.01 mg/ml Hoechst 33342 in PBS for 15 min at RT. After subsequent washes in PBS, the cells were mounted in Prolong gold antifade reagent and immediately imaged with a Zeiss Axio Observer Z1 microscope equipped with a 20x long distance Plan+Neofluar fluorescence objective (N/A=0.4 Ph2 Korr μ/1.5) and an AxioCam MRm fluorescence CCD camera. Each condition was performed in triplicates and 10 images per well were analyzed with AxioVision (Carl Zeiss) by measuring intensity of pixel^2^. The ratio between BODIPY^TM^ 493/503 and Hoechst 33342 was calculated and mean values normalized to control. Pooled results for repeated experiments are shown.

For visualization of LDs, GLUT1 and ZO-1, HUVECs grown for indicated time periods on 0.1% gelatin-coated glass chamber slides (4- or 8-well; BD Falcon) in FA-supplemented or control media were washed once in PBS and fixed for 10 min on ice with 4% PFA. After permeabilization for 30 min with PBS/0.2% BSA/0.1% saponin, cells were incubated with primary antibodies: guinea pig anti-adipophilin/PLIN2 (1:200; 20R-AP002, Fitzgerald Industries International), mouse anti-ZO-1 (1:500; 33-9100, Zymed) or rabbit anti-GLUT1 (1:200; 07-1401, Millipore) in PBS/0.2% BSA/0.1% saponin overnight at 4°C. Cells were thereafter washed in PBS/0.2% Tween-20 and Alexa Fluor® conjugated secondary antibodies and DAPI were applied for 1 h in PBS/0.2% BSA/0.1% saponin at room temperature. After subsequent washes in PBS/0.2% Tween-20, PBS and finally H_2_O, the cells were mounted in Prolong gold antifade reagent. Representative maximum intensity projections (2D renderings) of 4 μm confocal z-stacks from PLIN2 and ZO-1 immunostainings acquired with a Zeiss confocal laser scanning microscope (LSM700) equipped with a 63x Oil (N/A = 1.4) objective are presented. For quantification of GLUT1 signal using integrated density, each condition was performed in quadruplicates and five maximum intensity projections per well acquired with a confocal laser scanning microscope (LSM 700, Zeiss) equipped with 20x (N/A=0.8) objective and the ZEN 2009 software (Carl Zeiss Microimaging GmbH) using the same settings (within the respective experiment) were processed and analyzed using Adobe Photoshop and ImageJ. Mean values were normalized to control and pooled results for repeated experiments are shown.

### Trans-endothelial electrical resistance in primary human endothelial cells

For evaluation of cellular integrity, HUVECs were seeded onto gelatin-coated transwell inserts (BD Falcon) with 0.4 μm pore size suitable for 24-well plates. Endothelial growth medium (EGM-2) was added to the upper and lower chamber media and exchanged every day until a tight monolayer was formed. Thereafter, FA-supplemented (0.5 mM oleic acid) or control EGM-2 was added for 24 h. Trans-endothelial electrical resistance (TEER) was measured using a Milli-cell-ERS electrode from Millipore (Billerica, USA). Calibration of the electrode in reference media was performed before measurements and values from transwells without cells served as blank controls.

### Mice and diets

Animal work was approved by the Stockholm Animal Ethical Committee (ethical permits: 12790/23, 14960/20, 10627/18, 249/15, 187/13 and 650/12), or by the University of Michigan Institutional Animal Care and Use Committee (PRO00010905).

Animal experiments were conducted in accordance with Swedish animal welfare legislation and the European Union Directive (2010/63/EU) and performed in compliance with the guidelines from the Swedish National Board for Laboratory Animals or the United States Public Health Service’s Policy on Humane Care and Use of Laboratory Animals. The STAIR recommendations for preclinical development of acute ischemic stroke therapies were applied.^84^

For targeted deletion of the *Vegfb* allele, *loxP* sites were introduced in intron 1 and intron 6 of *Vegfb* (C57BL/6J NTac-*Vegfb*^tm3428Arte^; Taconic Biosciences, Germany). After arrival in the local animal facility, *Vegfb*^fl/+^ mice were backcrossed on C57BL/6J background (Charles River, Germany). Adipocyte-specific Cre expressor (Adipoq^Cre+^) mouse strain were obtained from the Jackson Laboratory and backcrossed on C57BL/6J background (Charles River). To generate mice with adipocyte-specific heterozygous deletion of VEGF-B (Adipoq^Cre+^;*Vegfb*^fl/*+*^mice), and relevant control genotypes, *Vegfb*^fl/+^ mice were bred with Adipoq^Cre+^ mice. Global *Vegfb* knock-out mice were generated by CRISPR-Cas9 mediated deletion in exon 3 resulting in a frame shift (obtained from Innovative Research, Novi, MI). Global *Vegfb* knock-out mice were backcrossed on C57BL/6J background (Charles River or Jackson Laboratory) before use.

Wild-type or genetically modified mice on C57BL/6J background (3-6 weeks old; Jackson Laboratory or Charles River) were randomly assigned a high-fat diet (HFD: 60 kcal% fat, 20 kcal%, protein and 20 kcal% carbohydrates; D12492, Research Diets) or standard grain-based chow diet (20 kcal% fat), maintained for 15-18 weeks until the experimental endpoint, when mice were around 20 weeks old. Mice were housed in groups of 4-5 in polystyrene cages with wooden chips as bedding and cellulose and paper as nesting material in a pathogen-free and climate-controlled environment with regulated 12 h light-dark cycles with *ad libitum* access to water and respective diet. Mice maintained on HFD develop diet-induced obesity, a condition characterized by hyperglycemia, dyslipidemia and insulin resistance, modeling obesity and type-2 diabetes ^21,36^. Body weights and blood glucose levels were monitored in all mice. Post-prandial blood glucose level was determined in the morning between 10 a.m. and noon after removing the food for 2 h. Blood glucose was measured using a hand-held glucose meter (Bayer AB) from a small drop of blood taken from the tail vein.

### In vivo antibody treatment

Age-matched wild-type chow or HFD mice were randomly assigned to treatment with neutralizing monoclonal anti-VEGF-B antibodies (400 μg/mouse of the murine 2H10, or the corresponding humanized CSL346, antibody; Proprietary to CSL Limited, Parkville, Australia) or isotype-matched control antibodies (400 μg/mouse Ctrl IgG; Proprietary to CSL Limited, Parkville, Australia;). The murine 2H10 antibody was used unless specified in the text. The treatment was initiated 7 days before MCAO (pre-treatment, in total three intraperitoneal (i.p.) injections at 7 days, 3 days and 1 day before MCAO), or as a therapeutic treatment with a single i.p. injection administered at 1 h post MCAO. Separate mouse cohorts received antibody pre-treatment for 7 days before blood sampling in conjunction with euthanasia, or as a single i.p. injection 4 h before intravenous injection with rtPA or vehicle control.

### Specificity of the anti-VEGF-B antibody

VEGF-B binding assay: A Nunc MaxiSorb plate (Thermo Scientific) was coated with 50 ng/well VEGF-A_11-109_, VEGF-B_10-108_ and PlGF_19-118_ (all three proteins cloned and produced at AMRAD)^85,86^ overnight at 4°C. The plate was blocked with 2% BSA in PBS, 50 µL/well, for 2 h at room temperature. The αVEGF-B antibody was serial diluted in Antibody Buffer (0.5% BSA in Tris-buffered saline, 0.05% Tween-20), transferred to the washed plate and incubated for 2 h at room temperature. The plate was washed and the secondary antibody, anti-mouse Ig affinity isolated from sheep and HRP conjugated (Chemicon), was applied at 0.5 µg/mL in Antibody Buffer, 50 µL/well, and incubated for 30 min at room temperature. After washing, the plate was developed with TMB-E (Millipore), 50 µL/well, and stopped with 25 µL/well 2 M phosphoric acid and the absorbance read at 450 nm.

### Photo-thrombotic middle cerebral artery occlusion (MCAO) model

MCAO was induced as described previously.^30^ Briefly, mice (matched for age and sex) were anesthetized with chloral hydrate (450 mg/kg) or isoflurane and placed securely under a dissecting microscope. The left MCA was exposed, and a laser Doppler flow probe (Type N, Transonic Systems) was placed on the surface of the cerebral cortex located 1.5 mm dorsal median from the bifurcation of the MCA. This location was then marked with a permanent waterproof pen, allowing repeated measurement. The probe was connected to a flow meter (Transonic model BLF21) and relative tissue perfusion units (TPU) were recorded with a continuous data acquisition program (Windaq, DATAQ). Rose Bengal (Fisher Scientific) was diluted to 10 mg/ml in Lactate Ringer’s and injected into the tail vein or retro-orbitally (50 mg/kg body weight). A 3.5-mW green light laser (540 nm, Melles Griot) was directed at the MCA from 6 cm distance, and the TPU of the cerebral cortex was recorded. Stable occlusion was achieved when the TPU dropped to less than 20% of pre-occlusion levels and did not rebound within 10 min after laser withdrawal. All cerebral blood flow (CBF) tracings were started 10 min before Rose Bengal injection and the average CBF over this time was considered 100% and used to normalize CBF. Time zero was set at Rose Bengal injection. CBF was then measured at 10 min and 30 min after MCAO. In a subset of mice, a milder photothrombotic insult was achieved by reducing the Rose Bengal dose (12.5 mg/kg body weight), and these animals were re-anesthetized at 24 h and 72 h after MCAO, and the Doppler flow probe was re-attached to the same location marked as before after the surgical site was re-exposed, to obtain 24 h and 72 h CBF data. Sham-operated mice were generated in parallel as negative controls. The sham mice were anesthetized and subjected to surgery and craniotomy but omitting Rose Bengal. Local (bupivacain) and systemic (5 mg/kg carprofen, occasionally buprenorphine) analgesia was administered for 24-48 h post-surgery.

Late thrombolysis with recombinant tPA was initiated 5 h post-occlusion in HFD mice treated with isotype control or αVEGF-B antibodies in a therapeutic setting (1 h post MCAO). Recombinant murine tPA (10 mg/kg body weight; Molecular Innovation, Novi, MI) was given as an intravenous bolus of 100 μl, followed by infusion with another 100 μl at 3 μl/min for 33 min.

For assessment of infarct volume, mice were sacrificed 72 h after MCAO and brains were removed and cut into 2 mm thick coronal sections and stained with 4% 2,3,5-triphenyltetrazolium chloride (TTC) in PBS for 20 min at 37°C, and then fixed in 4% paraformaldehyde solution for 10 min. Four brain slices/mouse were analyzed using the Image J software (NIH) by an investigator blind to the treatments. The following formula was used to calculate infarct volume.^30^

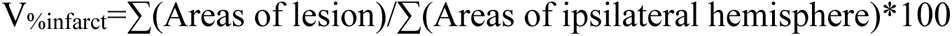

Infarct volume (V_%infarct_) was calculated as percent of the ipsilateral hemisphere in order to avoid an artifact due to brain edema.^87^

Intracerebral hemorrhage (ICH) was assessed at 72 h by a hemorrhage scoring system using the slices from the TTC staining. Scores; 0-4, where score 0 represent no signs of hemorrhage, score 1: 1-3 small petechial(s), score 2: 4-6 small petechials, score 3: >6 small petechials, and score 4: large area(s) of hemorrhage, were recorded and summed from 8 images per brain by an investigator blind to the treatments and average score/treatment group was thereafter calculated.

### Cerebral blood flow measurements

For full-field imaging of CBF, laser speckle contrast imaging was performed. Animals were anesthetized with isoflurane and scalp retracted to expose the intact cranium and then placed under a portable Laser Speckle device (MoorFLPI, Moor Instruments) connected to a computer equipped with real-time data acquisition software (MoorFLPI software version 2.01, Moor Instruments). A region of interest was defined (approximately 5 mm x 5 mm) over the MCA territory, and the mean flow in that region was measured before MCAO and then 3 h after MCAO.

### Thrombosis evaluation

FITC-conjugated murine fibrinogen (0.3 mg/kg) was injected intravenously via the tail vein 10 minutes before MCAO.^88^ Three hours after MCAO, mice were anesthetized with isoflurane and then perfused with PBS. Brains were harvested and embedded in OCT blocks. Frozen coronal sections were obtained, and infarct regions were assessed using an epi-fluorescence microscope (Eclipse TE200-E, Nikon). Images were then acquired using MetaMorph Imaging software (Molecular Devices, Sunnyvale, CA) and 5 fields of images obtained using the 40x objective were randomly chosen near the penumbra region and quantified using ImageJ software.

### Assessment of vascular leakage

For analysis of vascular leakage, mice were intravenously injected via the tail vein with 2.5 mg TMR-conjugated 70 kDa dextran (Dex^TMR^) in PBS (100μl; D1818, Molecular Probes) 1 h before sacrifice (or 10 min before MCAO if sacrificed within 2h). At 10 min, 0.5, 2, 3 or 5 h after MCAO, mice were thoroughly perfused with HBSS and 4% PFA for 4 min to ensure complete removal of intravascular tracer from the circulation. In a subset of mice, administration of FITC-conjugated fibrinogen was preceding MCAO and dextran injection. Dissected brains were briefly post-fixated in PFA for 1 h, thereafter washed in PBS for 24 h at 4°C, and finally submerged overnight in cryopreservation medium (30% sucrose) before embedding and freezing in cryo-compatible medium (NEG 50; Thermo Fisher Scientific). Prior to embedding, the dextran signal from intact brains (dorsal or lateral view) were acquired using a fluorescence dissecting stereomicroscope (Lumar V12; Zeiss). Coronal sections (12 μm cryo sections; 30 μm vibratome sections) from two different positions (bregma, +1 and bregma, -1) were analyzed using a Zeiss Axio Observer Z1 inverted microscope equipped with a 5x fluorescence objective (N/A=0.13 Korr μ/-) and an AxioCam MRm fluorescence CCD camera, and the ZEN 2009 software (Carl Zeiss Microimaging GmbH). The dextran signal from entire coronal sections were obtained using a motorized table and individual image tiles, taken with the 5x objective, stitched together by the software acquiring a single collate image per animal and bregma position. All images were acquired using the same settings and the area of pixels above a set threshold was determined using ImageJ.

In non-stroked mice, vascular leakage was assessed using Evans blue. Mice were intravenously injected with 100 μl of 4% Evans blue dye (Sigma-Aldrich) and allowed to circulate for 1 h before cardiac perfusion and sacrifice. Brains were removed and homogenized in N,N-dimethylformamide (Sigma-Aldrich) and centrifuged for 45 min at 25,000 rcf (Eppendorf centrifuge, model 5417R). The supernatants were collected, and quantitation of Evans blue extravasation performed using a spectrophotometer as previously described^30^ and determined from the formula:

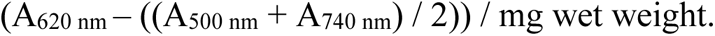

### In vivo rtPA treatment

Lyophilized rtPA (alteplase; Boehringer Ingelheim) was first dissolved in sterile MilliQ water and thereafter dialyzed against 0.4 M HEPES/0.1 M NaCl (pH7.4) using 1 kDa molecular weight cut-off dialysis tubes (Spectra/Por 6, Spectrum Laboratories) for long-term stability at -80°C. Concentration was determined using the A_280_ method with 1/E=1.9 correction. Ten mg rtPA per kg body weight was injected intravenously in the lateral tail vein as a bolus. Corresponding volume of vehicle (0.4 M HEPES/0.1 M NaCl) was injected to control mice.

### Immunohistochemistry

Non-stroked mice, or stroked mice at 3, 5, 5.5, 24 or 72 h after MCAO were terminally anesthetized and perfused transcardially with HBSS, followed by 4 % PFA. Dissected brains and livers (right lobe) were briefly post-fixated in 4 % PFA for 1 h and washed in PBS at 4°C for 24 h, and finally submerged in cryopreservation medium (30% sucrose O/N, 4°C) before embedding and freezing in NEG-50 (Thermo Fisher Scientific Inc.). Coronal cryo sections, 12 μm thick, were collected on Superfrost Plus slides (Thermo Fisher Scientific Inc.) and stored at -20°C until use. Slides were thawed, air-dried for 20 min at RT and washed in PBS to remove NEG-50. A permeabilization step for 10 min with 0.1% TritonX-100/PBS was performed before incubation for 1-2 h with serum-free protein block (X0909; Dako). For some targets: claudin-5, neuropilin-1 and VEGFR1, heat-induced epitope retrieval (S1699; Dako) in a kitchen steamer (1 h) was performed prior to permeabilization and block.

Visceral (epididymal and perirenal) and subcutaneous adipose tissue, and skeletal muscle (*m. quadriceps femoris*), were post-fixated O/N in 4% PFA, followed by several washes in PBS before dehydration and paraffin embedding. Four µm thick paraffin sections were mounted on Superfrost Plus slides. After rehydration, heat-induced epitope retrieval (S2367; Dako) in a kitchen steamer (1 h) was performed prior to permeabilization and block.

Primary antibodies were applied O/N at 4°C in blocking buffer: guinea pig anti-adipophilin/plin2 (1:200, 20R-AP002; Fitzgerald Industries International), goat anti-CD31 (1:200, AF3628; R&D Systems), rabbit anti-claudin-5 (1:200, 34-1600; Zymed), rabbit anti-Glut1 (1:200, 07-1401; Millipore), rabbit anti-hormone sensitive lipase (1:250, 4107; Cell Signaling), rabbit anti-phospho-serine_660_-hormone sensitive lipase (1:250, 45804; Cell Signaling), rabbit anti-Iba1 (1:200, 019-19741; Wako Chemicals), goat anti-neuropilin-1 (1:200, AF566; R&D Systems), goat anti-podocalyxin (1:200, AF1556; R&D Systems) and goat anti-VEGFR1 (1:200, AF471; R&D Systems). After several washes in PBS/0.1% Tween-20, Alexa Fluor® conjugated secondary antibodies were applied for 1 h at room temperature in blocking buffer: Donkey anti-goat or anti-rabbit IgG (Molecular Probes), or donkey anti-goat Fab or F(ab)_2_ antibodies (Jackson Immunoresearch). For plin2 immunostaining, a biotin-conjugated donkey anti-guinea pig IgG antibody (Jackson Immunoresearch) was used, followed by incubation with Alexa Fluor® conjugated streptavidin (Molecular Probes). Alternatively, a Cy-3 conjugated donkey anti-guinea pig antibody was used (Molecular Probes). Fab-specific secondary antibodies were used for detection of VEGFR1, and F(ab)_2_ antibodies were used for detection of neuropilin-1. For anatomical localization, nuclei were visualized with DAPI (4’,6-Diamidino-2-Phenylindole, Dihydrochloride, 0.2 μg/ml; Life Technologies). Finally, slides were mounted with cover glasses in Prolong Gold antifade reagent (P36930; Life Technologies).

All images were acquired with a confocal laser scanning microscope (LSM700; Zeiss) equipped with a 20x (N/A=0.8), 40x Oil (N/A =1.3) and 63x Oil (N/A = 1.4) objective, and the ZEN 2009 software (Carl Zeiss Microimaging GmbH). The images were processed and analyzed using Adobe Photoshop or ImageJ (National Institutes of Health). For quantification of antibody immunoreactivity using intensity, all images were acquired using the same settings (within the respective staining experiment) and the number of pixels above a set threshold was determined using the software. Each field of view analyzed was from a maximum intensity Z-stack image (10 μm stack) or from an epifluorescent image unless stated otherwise. The result from 6-9 fields of view (20 or 40x objective) from comparable anatomic positions in a given animal was averaged to obtain the value for that individual. Brightness and contrast settings were changed to generate final image and were applied equally to the entire image and within the same set of images. Adipocyte cell areas were measured in HSL-stained epididymal adipose tissue paraffin sections (all cells in 6 fields of view per mouse, 20x objective) using ImageJ. Images were evaluated by investigators blinded to study group. All immunostaining was repeated two – four independent times and representative images are shown.

### In situ RNA hybridization

For *in situ* RNA hybridization, mice were anesthetized with isoflurane and transcardially perfused with HBSS and 4% PFA at 3 h and 24 h after MCAO. Brains were post-fixated in PFA O/N, followed by several washes in PBS before dehydration and paraffin embedding. Five µm thick paraffin sections were subjected to RNA Scope® *in situ* hybridization protocol in strict accordance with the manufactureŕs instructions, including boiling the slides in Pretreat solution (RNA Scope® Vegfb target probe: Mm-Vegfb-CDS; Advanced Cell Diagnostics). The signal was developed using the 2.0 HD detection kit (brown chromogen; Advanced Cell Diagnostics) and sections lightly counterstained with Mayeŕs hematoxylin. Control slides and control probes (Advanced Cell Diagnostics) were run in parallel as positive and negative technical controls, and paraffin sections from naïve littermate *Vegfb*^+/+^ and *Vegfb*^-/-^ mice^89^ maintained on chow diet were used as biological positive and negative controls. Evaluation and image capture (63x objective) was done using a Zeiss Axio Observer Z1 inverted microscope and the ZEN 2009 software (Carl Zeiss Microimaging GmbH). Brightness and contrast settings were changed in Adobe Photoshop to generate final image and were applied equally to the entire image and within the same set of images. Representative images are shown.

### Transmission electron microscopy

Chow and HFD mice were subjected to MCAO. After 3 h, the mice were re-anesthetized with isoflurane and underwent transcardiac perfusion with 0.1 M Sorensen’s buffer followed by 4% paraformaldehyde and 2.5% glutaraldehyde in that buffer. Brains were then removed and an approximately 1.5 mm cube of cortical grey matter was taken from the ischemic penumbra, with the same location used for both chow and HFD mice and the same position also sampled in the contralateral hemisphere. Tissues were post-fixed in osmium tetroxide, dehydrated with ethanol, transitioned through propylene oxide and embedded in Epon. Ultrathin sections were examined on a Philips CM 100 electron microscope and images of the first 20 capillaries were recorded digitally with a Hamamatsu ORCA-HR camera system operated using AMT software (Advanced Microscopy Techniques Corp., Danvers, MA). The tissue processing, TEM and blood vessel imaging were performed by the Microscopy and Image Analysis Laboratory at the University of Michigan who were blinded to treatment. For each capillary profile, the distribution of LDs was examined and the number of endothelial LDs per profile determined. In addition, the dimensions (area, length and width) of each endothelial mitochondrion were determined using ImageJ and an average endothelial mitochondrion area for each sample calculated.

### Quantitative real-time PCR

Snap frozen ipsi- and contralateral brain hemispheres were homogenized using ceramic beads (PeriCellyse system, Bertin Technology) and RNA and protein simultaneously purified with the Nucleospin® TriPrep kit (Macherey-Nagel) according to the manufactureŕs instructions. cDNA synthesis was made using the iScript kit (Biorad) according to the manufactureŕs protocol. Quantitative real-time PCR (qPCR) was performed using primers for the genes: *Slc2a1* (GLUT1), *Vegfb*, *Flt1* (VEGFR1)*, Nrp1* and *Rpl19* as reference gene. SYBR green-based polymerase reaction using the SYBR^®^FAST kit (KAPA Biosystems) was performed in a Rotogene PCR machine (Qiagen) according to the manufactureŕs instructions. Expression values were calculated by the delta Ct-method and then converted to % of *Rpl19*, assuming a perfect primer-efficiency (2-fold product increase per cycle). Mouse primers used (5’ to 3’): m*Slc2a1* (GLUT1) forward: ATTGTGGCCGAGCTGTTC and reverse: GAGCACCGTGAAGATGATGA; m*Vegfb* forward: TCTGAGCATGGAACTCATGG, and reverse: TCTGCATTCACATTGGCTGT; m*Nrp1* forward: GGAGCTACTGGGCTGTGAAG and reverse: CCTCCTGTGAGCTGGAAGTC; m*Flt1* (VEGFR1) forward: GGAGGAGTACAACACCACGG and reverse: TTGAGGAGCTTTCACCGAAC; m*Rpl19* forward: GGTGACCTGGATGAGAAGGA and reverse: TTCAGCTTGTGGATGTGCTC.

### Blood lipid measurements

Mouse: EDTA-treated blood collected by cardiac puncture from terminally anesthetized mice or immediately after CO_2_ euthanasia was spun for 10 min at 2,000 x g in a cold centrifuge to obtain plasma. Plasma aliquots were snap frozen in aliquots at -80°C until further use. Commercially available kits were used for determination of triglycerides (Serum triglyceride determination kit; TR0100; Sigma), non-esterified fatty acids (NEFA-HR(2); Wako Chemicals), β-hydroxybutyrate (Stanbio Laboratories) and HDL and VLDL/LDL cholesterol (BioVision). Samples were run in 96-well format as duplicates together with a standard curve. The data was repeated at least twice, and by independent investigators blinded to study group.

Human: Pre-thrombolysis and matching post thrombolysis blood samples were obtained from 55 patients with acute ischemic stroke from a single stroke center biobank (Department of Neurology, University of Debrecen, Hungary). Blood was drawn within 1 h after the end of rtPA infusion, human ethics approval: RKEB/IKEB 3287-2010. Intravenous thrombolytic therapy was applied according to the European Stroke Organization (ESO) guidelines for rtPA (Alteplase, Boehringer Ingelheim, Germany) administration.^90^ Inclusion and exclusion criteria of patients were identical to standard criteria of thrombolysis as described in the ESO guidelines. Admission computed tomography angiography (CTA) imaging data was used to identify clot burden and location. None of the Hungarian patients were subjected to endovascular clot retrieval (ECR). Samples were collected between 2011 and 2013. Peripheral blood samples collected in sodium citrate tubes were processed immediately (centrifugation twice at 1500 g, RT, 15 min). Citrated plasma aliquots were stored deidentified at - 80°C. Comprehensive lipid profile including TGs was measured by standard laboratory methods using a Cobas 6000 automated analyser (Roche Diagnostics, Mannheim, Germany and Sysmex Europe GmbH, Hamburg, Germany). Plasma samples were sent to Australia and evaluated for non-esterified fatty acid (NEFA) levels using the NEFA-HR-2 assay (FujiFilm, product 434-91791, supplied by Wako Chemicals, GmbH, Germany) and tested at a 1:2 dilution. Human ethics approval was obtained from Monash University Human Research Ethics Committee: ID 21458.

### Ex vivo lipolysis

Visceral (epididymal and perirenal) and subcutaneous white adipose tissue were carefully dissected out immediately after HBSS perfusion of terminally anesthetized mice and cut into 20 mg pieces and put in prewarmed (37°C) DMEM (1 g/l glucose; Gibco) before use. The explants were transferred within 2 h to 200 μl DMEM containing 2% FA-free BSA (Sigma-Aldrich) and incubated for 1 h at 37°C, 5% CO_2_, in the presence or absence of 10 μM isoproterenol (Cayman Chemical) to measure stimulated lipolysis. Thereafter, the media was collected and NEFA content was determined using commercially available kits for enzymatic determination of NEFAs (Wako Chemicals). Total protein content (BCA protein assay; Pierce) in the explants was measured and used for normalization. Tissue explant lysates were prepared after extraction of lipids with chloroform/methanol (2:1, v/v), 1% glacial acetic acid, for 1 h, 37°C under vigorous shaking, followed by lysis in 0.3 N NaOH/0.1% sodium dodecyl sulphate O/N at 55°C under vigorous shaking. NEFA release was analyzed from two explants per fat pad and experimental condition, and averaged. *Ex vivo* lipolysis rate was calculated as nmol NEFAs released per h and mg protein content.

### Positron emission tomography

Positron emission tomography (PET) was performed on a Focus120 MicroPET (CTI Concorde Microsystems). PET data were processed with MicroPET Manager (CTI Concorde Microsystems) and evaluated using the Inveon Research Workplace software (Siemens Medical Solutions). HFD or chow fed age-matched mice were anaesthetized with isoflurane (5% initially and 1.5% to maintain anesthesia) and placed on the camera bed on a heating pad (37°C). 2-[^18^F] fluoro-2-deoxy-D-glucose, ^18^F-DG, was synthesized using an automated synthesis module (Fastlab, General Electric Medical Systems AB) and had passed routine clinical quality controls according to European Pharmacopoeia, Ph. Eur. Mice received tail vein intravenous injections of 10 MBq ^18^F-DG in a volume of 200 μl. The PET data acquisition started at injection. All PET data were normalized with respect to the injected dose of each mouse (% ID/g). The signal was collected from a region of interest of 30-35 mm^3^ including the hippocampus.

### Statistical analysis

All data represent the mean ± s.e.m. *n* indicates the number of individual mice or tissue culture wells used in the study. No mice were excluded from the analyses. Statistical analysis was performed using GraphPad Prism 10 statistical software. Prior to statistical analysis, conformity to normality distribution of the datasets were determined. When applicable, conformity of data to log^10^ normality was applied before analysis. For comparison between two groups, two-tailed unpaired Student’s *t*-test was used. For multiple comparisons, one-way or two-way analysis of variance, ANOVA, followed by a multiple comparisons test, was utilized as indicated in the figure legends. When applicable, the appropriate non-parametric tests were used as indicated in the figure legends. A *P*-value less than 0.05 were considered statistically significant and indicated in the figures by asterisks.

