## Supplemental information for "Thrombolysis exacerbates cerebrovascular injury after ischemic stroke via a VEGF-B dependent effect on adipose lipolysis"

Supplementary Figure 1. Nilsson & Su et al. Related to Fig. 1

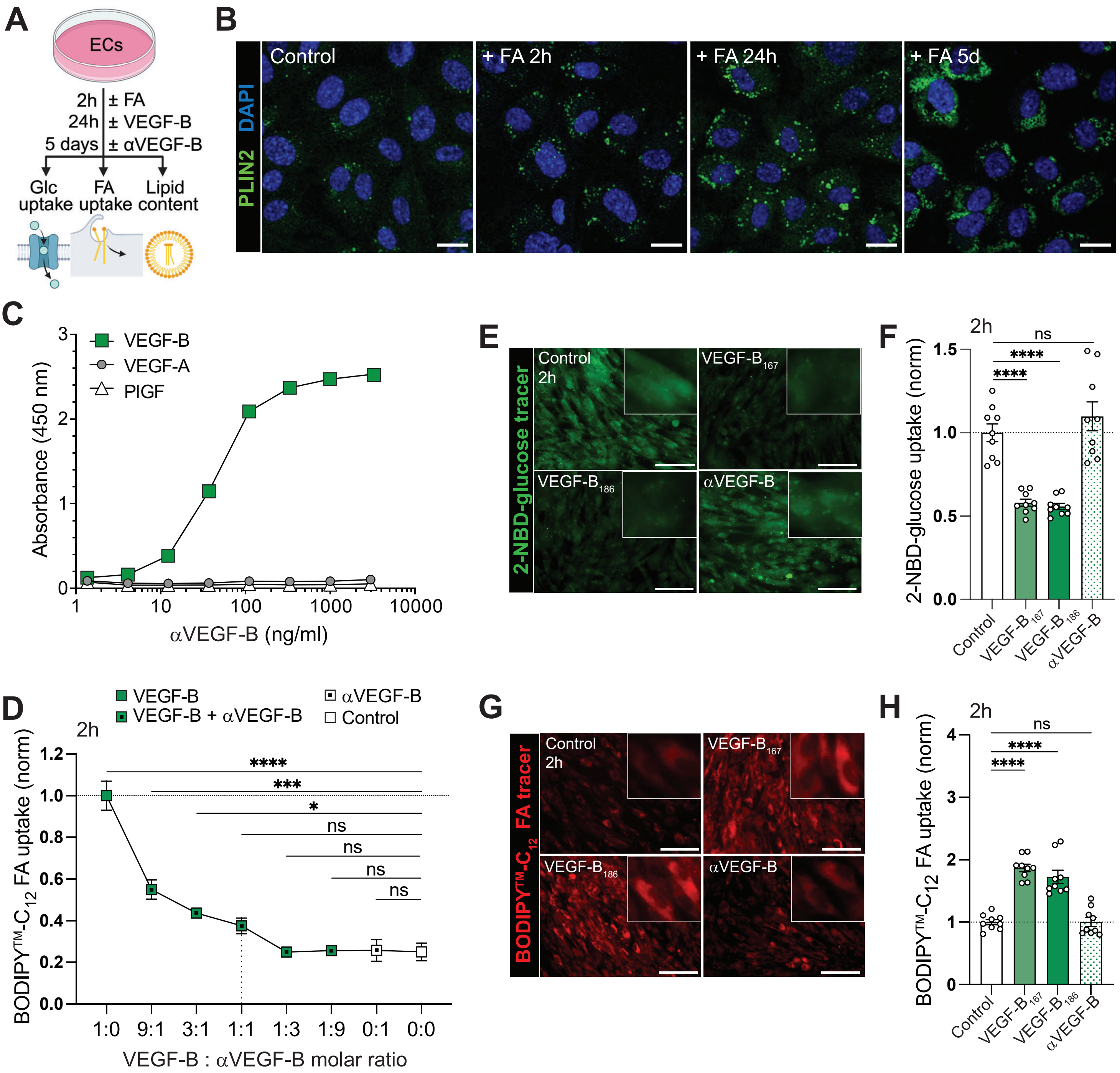

**Figure S1. Paracrine VEGF-B signaling induces a shift from glucose to fatty acid uptake in primary human brain endothelial cells (related to Figure 1). (A)** Schematic representation of experimental outline. **(B)** PLIN2+ LDs in primary human endothelial cells (HUVECs) exposed to 0.05 mM oleic/palmitic acid mix for 2 h, 24 h and 5 days. **(C)** Binding affinity and specificity of a neutralizing αVEGF-B monoclonal antibody to VEGF-B, VEGF-A or PlGF. **(D)** Neutralizing capacity of the αVEGF-B antibody on VEGF-B-induced long-chain FA uptake in HUVECs. **(E-H)** Glucose (E-F) and FA (G-H) uptake in primary human brain microvascular endothelial cells (HBMECs) after stimulation with recombinant VEGF-B protein (VEGF-$B_{167}$ or VEGF-$B_{186}$ isoforms) or the neutralizing αVEGF-B monoclonal antibody for 2 h. Insets show higher magnification (E, G). Quantification (F, H). Scale bars, 10 µm (B); 50 µm (E, G). Data presented as mean ± s.e.m of three (D) or nine (F, H) biological replicates per condition, normalized to VEGF-B (D) or control (F, H) treated cells. Statistical evaluation compared to control using ordinary one-way ANOVA (D, H) or Brown-Forsythe and Welch ANOVA (F), followed by Dunnett´s multiple comparisons test. *$P$<0.05, ***$P$<0.001, ****$P$<0.0001.

Supplementary Figure 2. Nilsson & Su et al. Related to Fig. 2

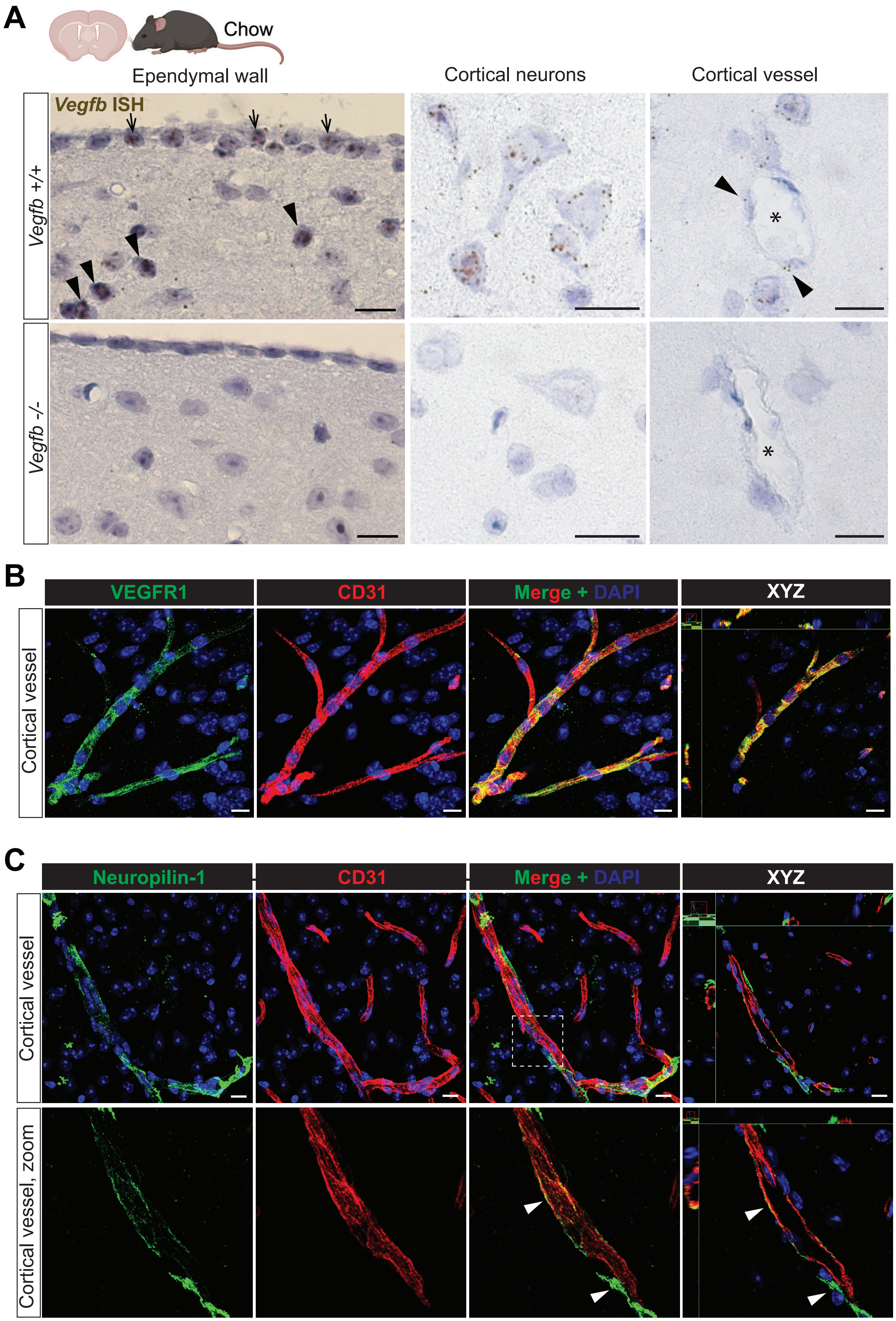

**Figure S2. VEGF-B and VEGF-B receptor expression in murine brain (related to Figure 2). (A)** *Vegfb* mRNA *in situ* hybridization using RNA Scope technology (brown dots). Counterstaining with hematoxylin in blue. *Vegfb* expression in cells lining the ventricle walls marked with arrows (upper left panel). *Vegfb* expression in periventricular neurons (upper middle panel) and perivascular cells (upper right panel, arrowheads). Blood vessel lumen indicated with asterisks (right panels). *Vegfb*-deficient mice (*Vegfb*$^{-/-}$) included for probe specificity (lower panels). **(B-C)** VEGFR1 (B), the main receptor for VEGF-B, and the co-receptor neuropilin-1 (C) expression in brain. Co-localization with CD31 is shown. Neuropilin-1 expression in perivascular cells (lower arrowhead in C). Scale bars, 10 μm.

Supplementary Figure 3. Nilsson & Su et al. Related to Fig. 2 and 3

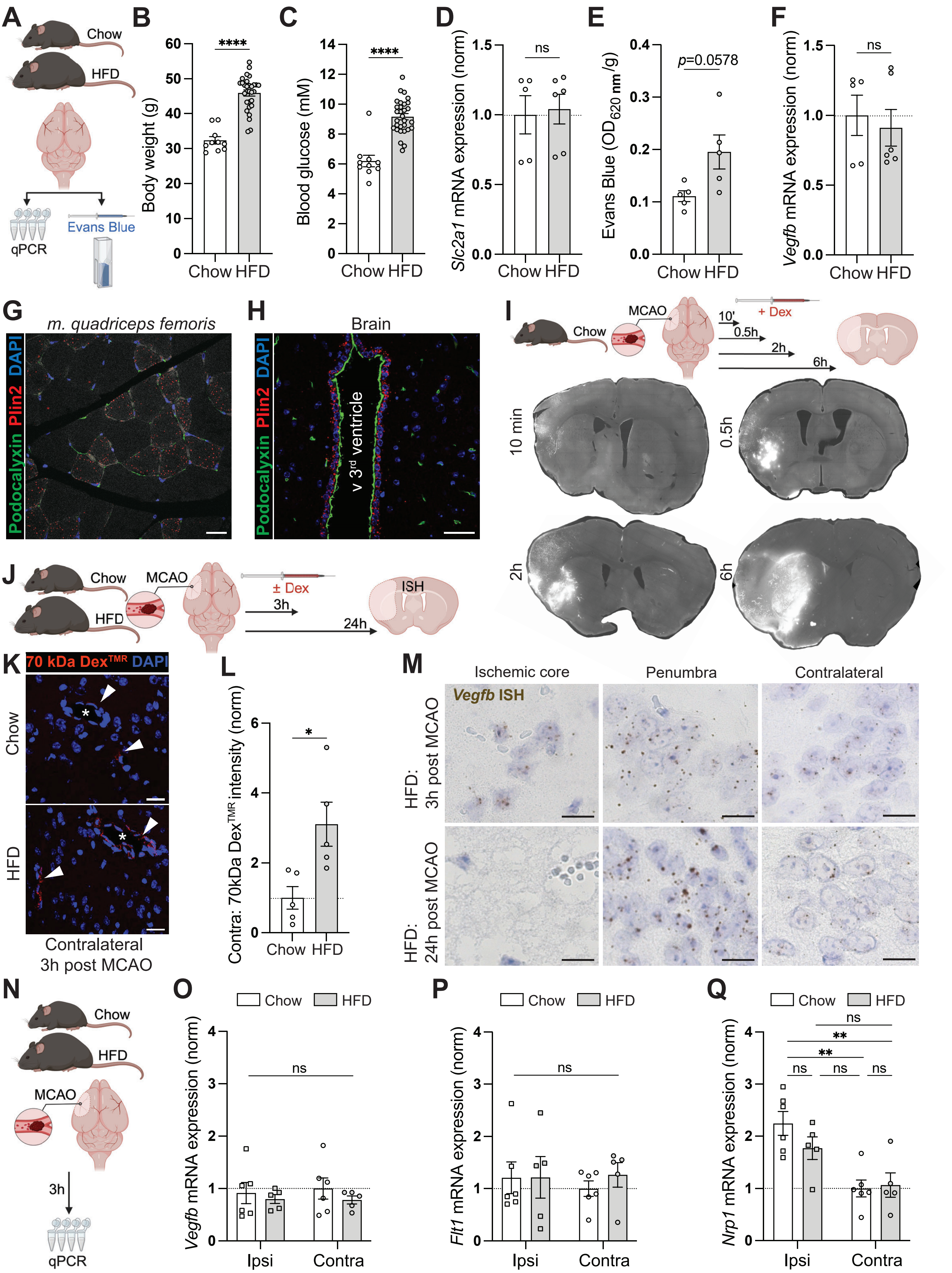

**Figure S3. Vascular permeability progression and effect of HFD on expression of VEGF-B and its receptors in brain after acute ischemic stroke (related to Figure 2 and 3). (A)** Schematic representation of experimental outline. HFD mice compared to age-matched littermate control mice maintained on grain-based standard chow. **(B)** Body weight. (**C)** Blood glucose. **(D)** Brain *Slc2a1* (Glut1) mRNA expression. **(E)** Evans Blue permeability in brain. **(F)** Brain *Vegfb* mRNA expression. **(G)** Plin2+ LDs in a mixed fiber skeletal muscle biopsy (*m. quadriceps femoris*) from a chow-fed mouse. Endothelial cell co-staining with podocalyxin. **(H)** Plin2+ LDs in the ependymal wall, ventral 3$^{rd}$ ventricle. Endothelial cell co-staining with podocalyxin, also marking ependymal cell glycocalyx. **(I)** Dex$^{TMR}$ extravasation at 10 min, 0.5 h, 2 h and 6 h post MCAO (50 mg/kg Rose Bengal) in chow-fed mice. Dex$^{TMR}$ injected intravenously 10 min before MCAO in mice analyzed up to 2 h post MCAO and 1 h before sacrifice in mice analyzed at 6 h post-MCAO. **(J)** Schematic representation of experimental outline. **(K-L)** Age-matched chow or HFD mice subjected to MCAO (50 mg/kg Rose Bengal) and sacrificed 3 h later. Dex$^{TMR}$ tracer in contralateral (non-ischemic) hemisphere. Blood vessel lumen marked with asterisks (K). Quantification of contralateral Dex$^{TMR}$ signal intensity (L). **(M)** *Vegfb* mRNA *in situ* hybridization (brown) 3 h and 24 h post MCAO (HFD mice). Counterstaining with hematoxylin (blue). (**N)** qPCR analyses on RNA purified from whole ischemic (Ipsi) and contralateral (Contra) hemispheres 3 h after MCAO in chow and HFD mice. **(O-Q)** Expression of *Vegfb* (O), *Flt1* (VEGFR1) (P) and *Nrp1* (neuropilin-1) (Q). Scale bars, 50 µm (G-H); 20 µm (K); 10 µm (M). Data presented as mean ± s.e.m of n=10 (chow), n=30 (HFD) mice/group (B-C) and n=5-6 mice/group (D-F, L, O-Q). Statistical evaluation using unpaired *t*-test (B-C, F, L), Mann-Whitney test (D), unpaired *t*-test with Welch´s correction (E) or two-way ANOVA followed by Tukey´s multiple comparisons test (O-Q). *$P<0.05$, **$P<0.01$, ****$P<0.0001$.

Supplementary Figure 4. Nilsson & Su et al. Related to Fig. 4

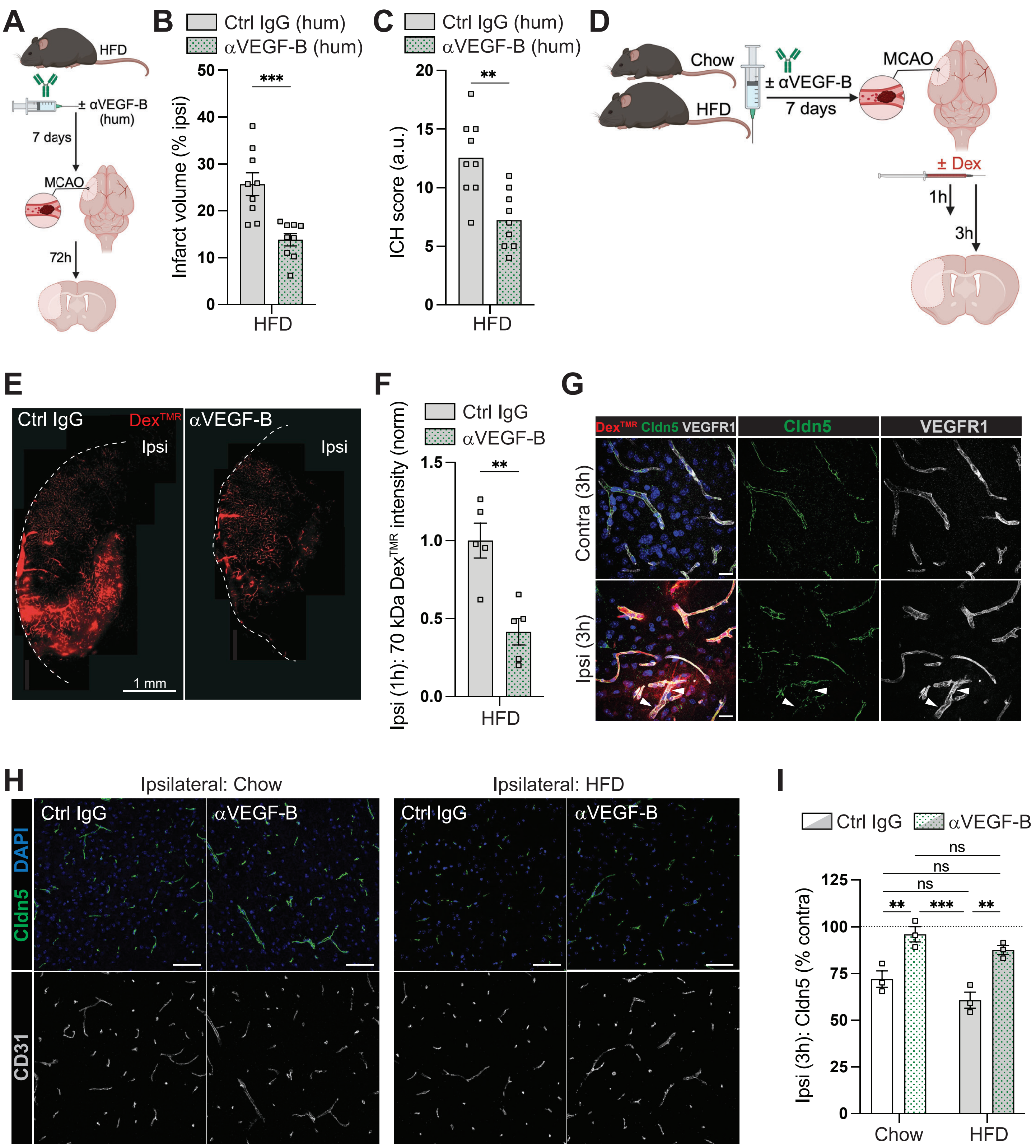

**Figure S4. Pre-treatment with neutralizing anti-VEGF-B antibodies improves blood-brain barrier integrity in HFD mice (related to Figure 4). (A)** Schematic representation of experimental outline. HFD mice pre-treated for 1 week with humanized isotype control (Ctrl IgG hum) or humanized neutralizing αVEGF-B antibodies (αVEGF-B hum) before subjected to MCAO (50 mg/kg Rose Bengal) and sacrificed 72 h later. **(B)** Infarct volumes. **(C)** ICH scores. **(D)** Schematic representation of experimental outline. Chow or HFD mice pre-treated for 1 week with isotype control (Ctrl IgG) or αVEGF-B antibodies before subjected to MCAO (50 mg/kg Rose Bengal). **(E-F)** $Dex^{TMR}$ distribution (E) and quantification (F) in HFD mice 1 h post-MCAO ($Dex^{TMR}$ injected intravenously 10 min before MCAO). **(G)** Claudin-5 intensity in permeable blood vessels in chow fed mice 3 h post MCAO. Arrowheads point to VEGFR1+ vessel portions that have lost claudin-5. **(H-I)** Claudin-5 distribution (H) and quantification (I) in CD31+ vessels 3 h post MCAO. Scale bars, 1 mm (E); 20 µm (G); 50 µm (H). Data presented as mean ± s.e.m of n=9 mice/group (B-C), n=5 mice/group (F) and n=3 mice/group (I). Statistical evaluation using unpaired *t*-test (B-C, F), or two-way ANOVA followed by Tukey´s multiple comparisons test (I). $^{**}P<0.01$, $^{***}P<0.001$.

Supplementary Figure 5. Nilsson & Su et al. Related to Fig. 4

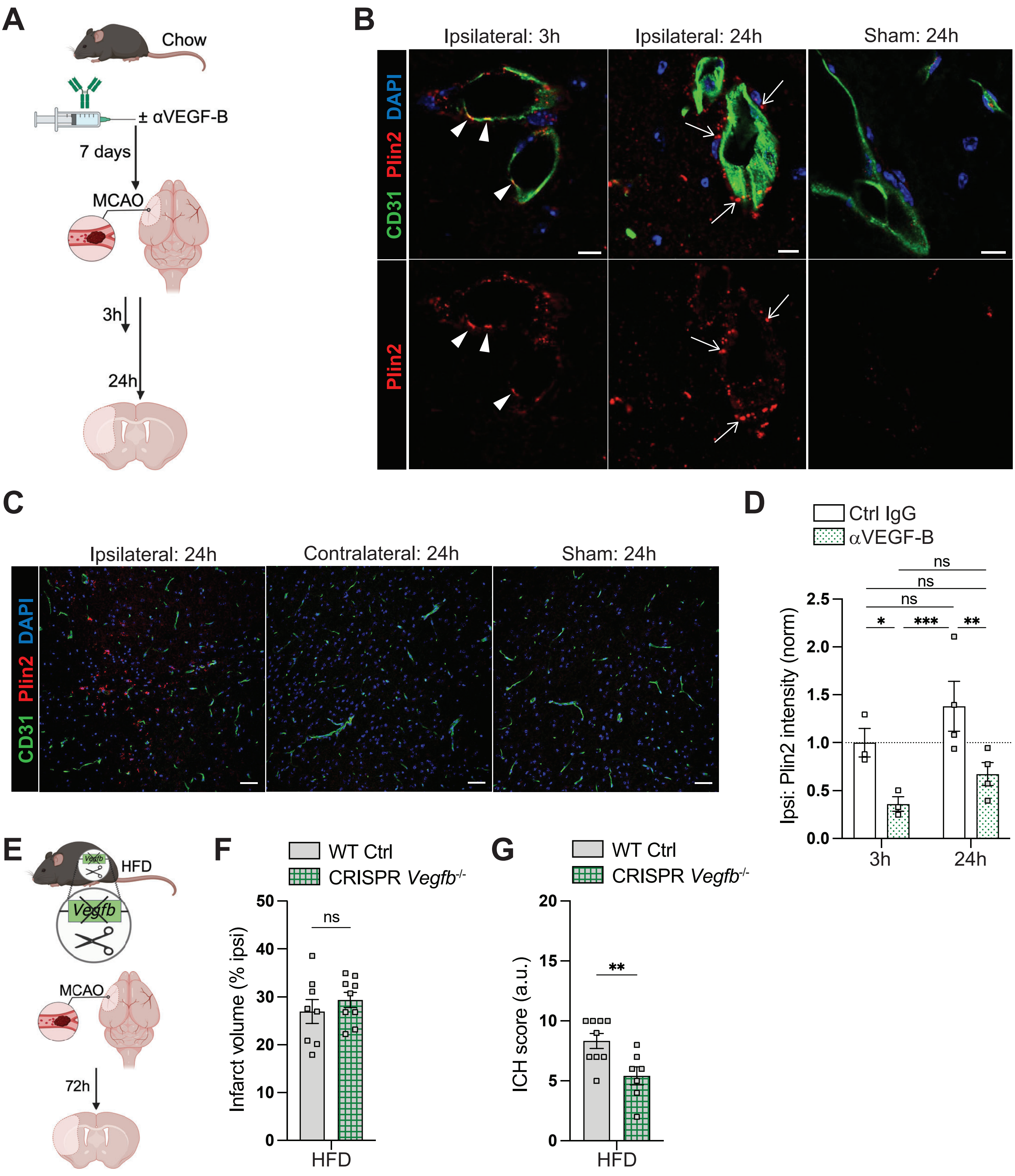

**Figure S5. Effect of pre-treatment with neutralizing anti-VEGF-B antibodies and congenital *Vegfb* deficiency on outcome after ischemic stroke (related to Figure 4). (A)** Schematic representation of experimental outline. Chow-fed mice pre-treated for 1 week with isotype control (Ctrl IgG) or αVEGF-B antibodies (αVEGF-B) before subjected to MCAO (50 mg/kg Rose Bengal) and sacrificed 3 h or 24 h later. Sham mice included as control for surgery and post-op procedures. **(B)** High magnification images showing Plin2+ LDs inside CD31+ endothelial cells (arrowheads) at 3 h post MCAO. At 24 h post MCAO, LDs are predominantly accumulating inside perivascular cells (arrows). **(C)** Overview images showing accumulation of Plin2+ LDs in dispersed brain parenchymal cells 24 h post-MCAO. **(D)** Quantification of Plin2 intensity. **(E)** *Vegfb* knock-out mice (*Vegfb*$^{-/-}$) created with CRISPR-Cas9 technology and *Vegfb*$^{+/+}$ littermate controls on HFD subjected to MCAO (50 mg/kg Rose Bengal) and sacrificed 72 h later. **(F)** Infarct volumes. **(G)** ICH scores. Scale bars, 10 µm (B); 50 µm (C). Data presented as mean ± s.e.m of n=3-4 mice/group (D) and n=8-9 (F-G) mice/group. Statistical evaluation using two-way ANOVA followed by Tukey´s multiple comparisons test (D) or unpaired *t*-test (F-G). **P*<0.05, ***P*<0.01, ****P*<0.001.

Supplementary Figure 6. Nilsson & Su et al. Related to Fig. 5

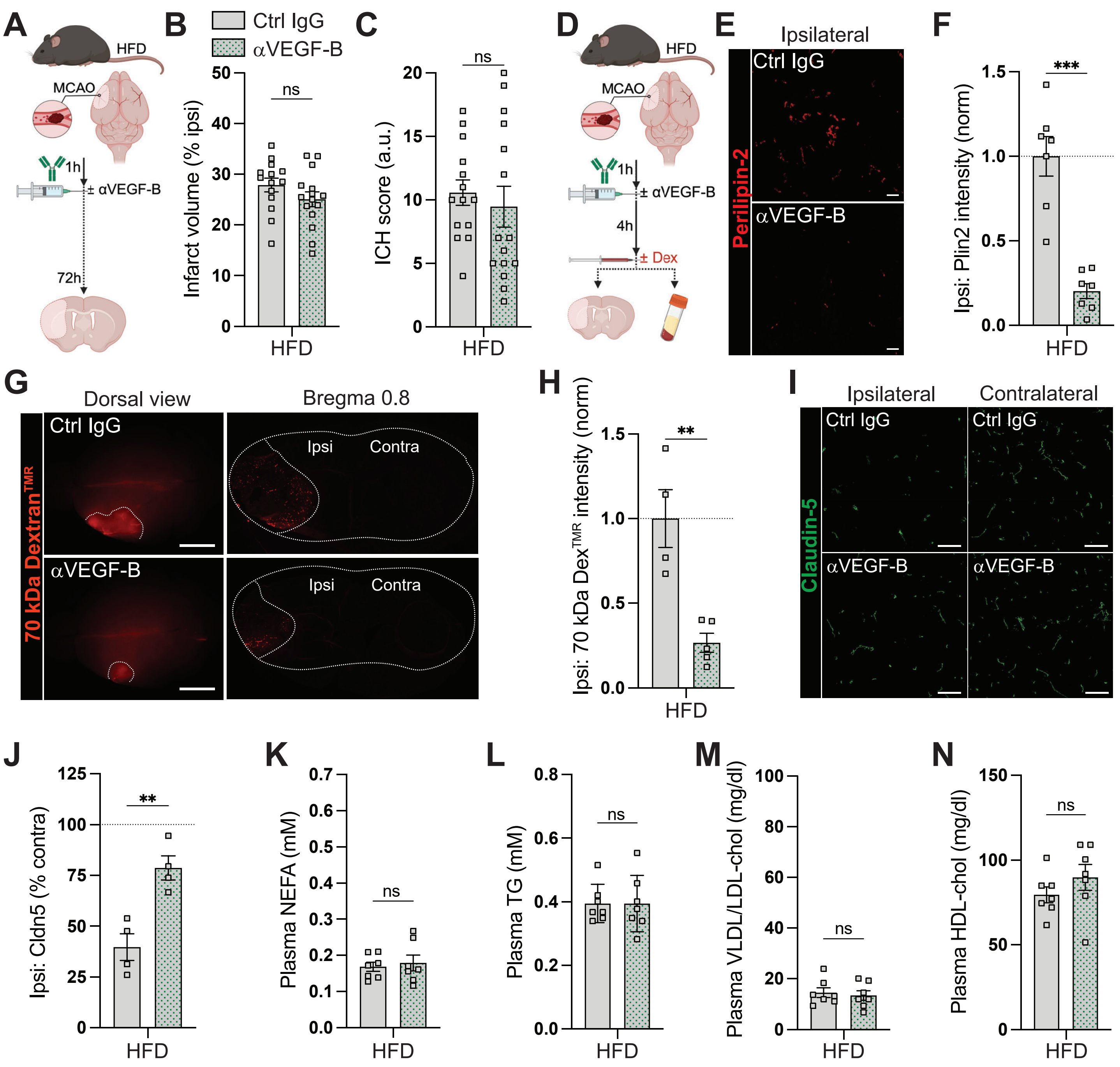

**Figure S6. Therapeutic treatment with neutralizing anti-VEGF-B antibodies improves blood-brain barrier integrity, correlating with decreased cerebrovascular lipid droplet accumulation (related to Figure 5). (A)** Schematic representation of experimental setup. HFD mice subjected to MCAO (50 mg/kg Rose Bengal) and 1 h later given one i.p. injection with isotype control (Ctrl IgG) or αVEGF-B antibodies. Experimental endpoint at 72 h post MCAO. **(B)** Infarct volumes. **(C)** ICH scores. **(D)** Schematic representation of experimental setup. HFD mice subjected to MCAO (50 mg/kg Rose Bengal) and 1 h later given one i.p. injection with isotype control (Ctrl IgG) or αVEGF-B antibodies. Experimental endpoint at 5 h post MCAO. **(E-F)**. Distribution (E) and quantification (F) of Plin-2+ LDs. **(G)** Dorsal whole-mount view and histological cross-sections showing $Dex^{TMR}$ tracer extravasation. **(H)** Ipsilateral $Dex^{TMR}$ signal intensity. **(I-J)** Claudin-5 distribution (I) and quantification (J). **(K-N)** Plasma NEFAs (K), TGs (L), VLDL/LDL cholesterol (M) or HDL cholesterol (N) levels. Scale bars, 5 mm (G); 50 µm (E, I). Data presented as mean ± s.e.m. based on n=14+15 (Ctrl IgG *vs.* αVEGF-B) mice/group (B-C), n=7 mice/group (F, K-N) and n=4-5 mice/group (H, J). Statistical evaluation using unpaired *t*-test or unpaired *t*-test with Welch´s correction (F). ***P*<0.01, ****P*<0.001.

Supplementary Figure 7. Nilsson & Su et al. Related to Fig. 6 and 7

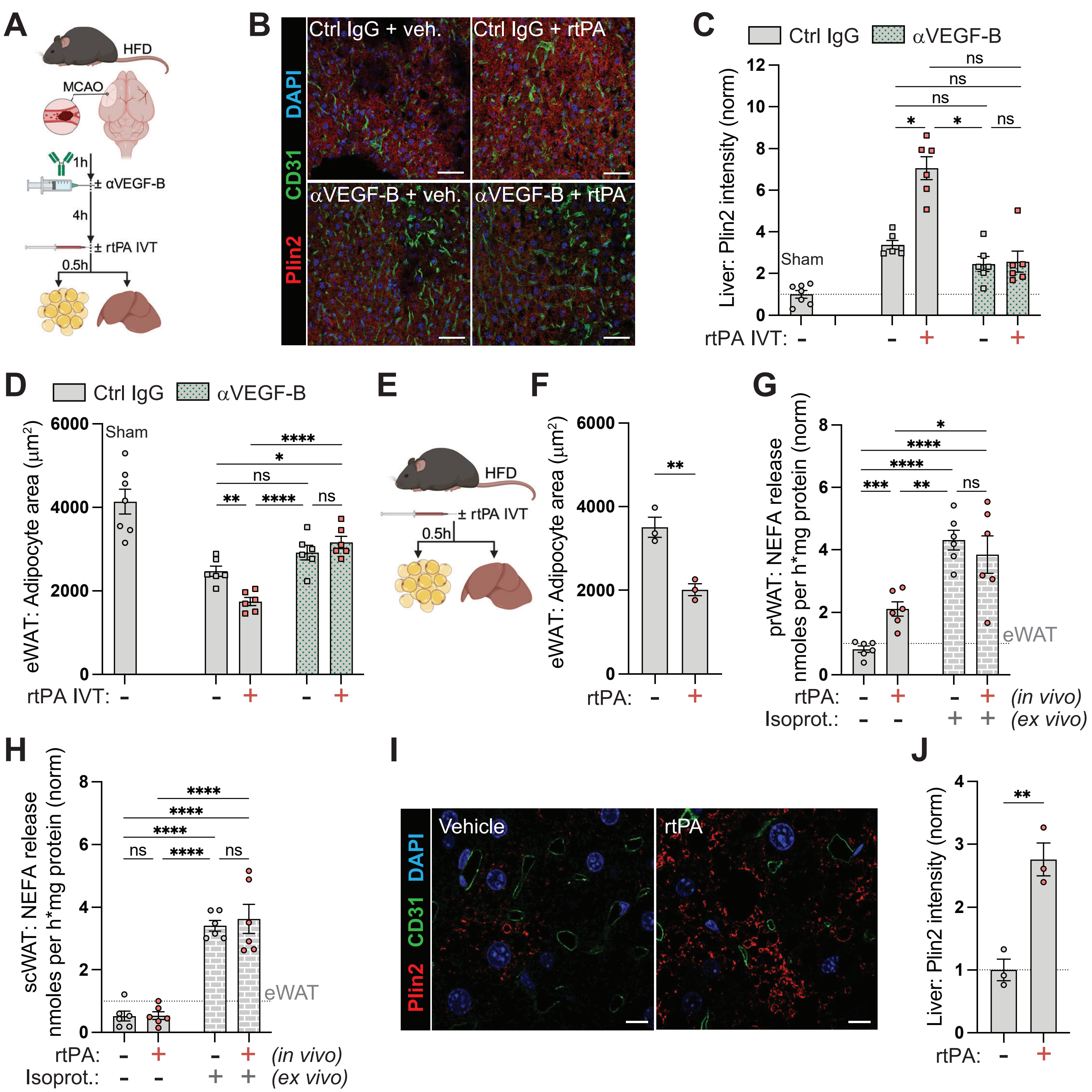

**Figure S7. Intravenous rtPA administration induces visceral adipose tissue lipolysis and promotes lipid droplet formation in liver (related to Figure 6 and 7). (A)** Schematic representation of experimental setup. HFD mice subjected to MCAO (50 mg/kg Rose Bengal) and 1 h later given one i.p. injection with isotype control (Ctrl IgG) or αVEGF-B antibodies, followed by intravenous thrombolysis (IVT) with rtPA (alteplase, 10 mg/kg) or vehicle as control, 4 hours later (5 h after MCAO). Experimental endpoint at 5.5 h post MCAO. Sham mice included as control for surgery and post-op procedures. **(B-C)** Plin2+ LDs in liver. Endothelial cells stained with CD31 (B). Plin2 quantification (C). **(D)** Adipocyte cell area in eWAT. **(E)** Schematic representation of experimental setup. Intravenous rtPA (alteplase, 10 mg/kg) injection of age-matched HFD mice. Experimental endpoint 30 min post rtPA injection. **(F)** Adipocyte cell area in eWAT. **(G-H)** *Ex vivo* lipolysis in perirenal (G) and subcutaneous (H) WAT (basal and isoproterenol stimulated). Values normalized to eWAT. **(I-J)** Quantification (I) and distribution (J) of Plin-2+ LDs in liver. Endothelial cells stained with CD31. **(**Scale bars, 50 μm (B); 10 μm (J). Data are presented as mean ± s.e.m of n=6-7 mice/group (C-D, G-H) and n=3 mice/group (F, I). Statistical evaluation using two-way ANOVA followed by Tukey´s multiple comparisons test (C-D, G-H) or unpaired *t*-test (F, I). **P*<0.05, ***P*<0.01, ****P*<0.001, *****P*<0.0001.
